# Towards the molecular architecture of the peroxisomal receptor docking complex

**DOI:** 10.1101/854497

**Authors:** Pascal Lill, Tobias Hansen, Daniel Wendscheck, Bjoern Udo Klink, Tomasz Jeziorek, Jonas Miehling, Julian Bender, Friedel Drepper, Wolfgang Girzalsky, Bettina Warscheid, Ralf Erdmann, Christos Gatsogiannis

**Affiliations:** Department of Structural Biochemistry, Max Planck Institute of Molecular Physiology, Dortmund, Germany; Institute of Biochemistry and Pathobiochemistry, Faculty of Medicine, System Biochemistry, Ruhr-University Bochum, Bochum, Germany; Biochemistry and Functional Proteomics, Institute of Biology II, Faculty of Biology, University of Freiburg, 79104 Freiburg, Germany; Signalling Research Centres BIOSS and CIBSS, University of Freiburg; Interdisciplinary Research Center HALOmem, Institute for Biochemistry and Biotechnology, Martin Luther University Halle-Wittenberg, Halle (Saale), Germany

**Author notes:** These authors contributed equally to this work (Pascal Lill, Tobias Hansen, Daniel Wendscheck). These authors contributed equally to this work (Bettina Warscheid, Ralf Erdmann).

## Abstract

Import of yeast peroxisomal matrix proteins is initiated by cytosolic receptors, which specifically recognize and bind the respective cargo proteins. At the peroxisomal membrane, the cargo-loaded receptor interacts with the docking protein Pex14p that is tightly associated with Pex17p. Previous data suggest that this interaction triggers the formation of an import pore for further translocation of the cargo. The mechanistic principles are however unclear, mainly because structures of higher order assemblies are still lacking. Here, using an integrative approach, we provide the first structural characterization of the major components of the peroxisomal docking complex Pex14p/Pex17p, in a native bilayer environment and reveal its subunit organization. Our data show that three copies of Pex14p and a single copy of Pex17p assemble to form a 20 nm rod-like particle. The different subunits are arranged in a parallel manner, showing interactions along their complete sequences and providing receptor binding-sites on both membrane sides. The long rod facing the cytosol is mainly formed by the predicted coiled-coil domains of Pex14p and Pex17p, possibly providing the necessary structural support for the formation of the import pore. Further implications of Pex14p/Pex17p for formation of the peroxisomal translocon are discussed.

## Introduction

Peroxisomes are organelles present nearly ubiquitously in eukaryotic cells, ranging from unicellular yeasts to multicellular organisms, such as plants and humans. Beside ß-oxidation of fatty acids as a main conserved function of peroxisomes, a broad range of additional metabolic functions is linked to this organelle, underscored by severe and frequently lethal phenotypes of human disorders(*1*, *2*). These organelles do not contain DNA and thus all peroxisomal matrix proteins are encoded in the nucleus and synthesized on free polyribosomes in the cytosol. Subsequently, matrix proteins are targeted to the organelle by peroxisomal import receptors(*3*). A remarkable feature of peroxisomes is that unlike the transport of unfolded polypeptides across the membranes of the endoplasmic reticulum, mitochondria, and chloroplast, they can import already folded, cofactor-bound, and even oligomeric proteins(*4*, *5*). This transport is highly selective and mediated by specific import sequences known as peroxisomal targeting signals (PTSs)(*6*, *7*). Peroxisomal matrix proteins equipped with either a carboxy-terminal PTS1 or an amino-terminal PTS2, are recognized and bound in the cytosol by the import receptor Pex5p or Pex7p, respectively(*8*, *9*). A peroxisomal membrane associated complex consisting of Pex13p, Pex14p and Pex17p in yeast allows docking of the cargo-loaded receptor(*10*–*14*). This primary interaction of the cargo-loaded receptor with the docking complex induces the formation of a transient and highly dynamic import pore, necessary for the translocation of the cargo across the peroxisomal membrane(*15*–*17*). How translocation and release of the cargo are realized in detail still remains enigmatic but it has been previously shown that the receptor is exported from the peroxisomal membrane in an ubiquitin- and ATP-dependent manner, a process that is discussed to provide the driving force for cargo-import according to the export-driven-import model(*18*–*20*).

The receptor-docking complex is of major importance for peroxisomal matrix protein import, as it provides a binding platform for newly formed receptor-cargo complexes at the peroxisomal membrane. Both Pex13p and Pex14p are peroxisomal membrane proteins providing several binding sites for the import receptors Pex5p and Pex7p(*16*). Docking of the Pex5p-PTS1 protein complex at the peroxisome membrane is supposed to occur at Pex14p(*21*, *22*). Pex17p is tightly associated with Pex14p(*23*), but its precise function remains unknown. Although Pex17p is part of the docking complex in yeast, it does not significantly contribute to the assembly of the Pex13p/Pex14p subcomplex(*15*, *23*, *24*), and its counterpart in higher eukaryotes has not yet been identified. However, Pex17p is essential for peroxisomal import of both PTS1 and PTS2 proteins(*14*). Strikingly, both import receptors, Pex5p and Pex7p, associate with the docking-complex (Pex13p, Pex14p) in absence of Pex17p, but with decreased efficiency(*24*). Pex17p is apparently non-essential for receptor docking, but it increases the efficiency of the docking event.

Furthermore, albeit a close association between the core components of the docking complex (Pex13p, Pex14p) is important for matrix protein import (*25*), there are several lines of evidence that Pex13p is not a permanent component of the peroxisomal docking complex or the import pore(*10*, *26*) and interestingly, an assembly between the receptor Pex5p and the docking component Pex14p in absence of Pex13p is capable per se to form a large transient channel at the peroxisome membrane(*15*).

However, little is known about the molecular mechanism underlying the primary docking and subsequent translocation events, largely because structures of the higher-order assemblies are not available. Here, using cryo-electron microscopy single particle analysis (cryo-EM SPA) and cryo-electron tomography (cryo-ET) combined with crosslinking and native mass spectrometry (MS), we set out to characterize the overall architecture of the yeast Pex14p/Pex17p complex.

## Results

### Pex14p forms a 3:1 heterotetrameric complex with Pex17p

Yeast Pex14p consists of three structural regions: a highly conserved N-terminal region containing a putative transmembrane domain, a long and a short coiled-coil domain in the middle region and an unstructured C-terminal domain (Figure 1a). Yeast Pex14p contains two separated binding sites for the PTS1 import receptor Pex5p, localized at its N- and C-termini (Figure 1a)(*27*). Mammalian PEX14 contains only one PEX5 binding site and only a single, shortened central predicted coiled-coil domain(*28*), which has been shown to mediate dimerization, whereas the transmembrane domain is responsible for formation of higher order assemblies(*29*). The topology of Pex14p at the peroxisomal membrane has been controversial, but recent results indicate that the N-terminus faces the matrix of the organelle(*30*). Pex17p contains a putative transmembrane domain near the N-terminus and two short coiled-coil domains (Figure 1a). Pex14p and Pex17p associate together to form a tight core complex, however the oligomeric composition of the complex is yet unknown. Interestingly in rice blast fungus *Mangaporthe*, Pex14 and Pex17 fuse to form a unique Pex14/Pex17 (or Pex33) peroxin(*31*). Here, we coexpressed yeast Strep_II_-Pex14p and His_6_-Pex17p in *E. coli* (see Material and Methods). Initial experiments revealed a dominant Pex14p-degradation product giving rise to a band in SDS-PAGE at about 35 kDa, which was most likely a consequence of a site-specific cleavage around the methionine at position 51, as identified by MS (Supplementary Figures 1a-c). Hence, we replaced methionine 51 by leucine and co-expressed this Pex14p-variant (Strep_II_-Pex14p(M51L)), for which the additional band at 35 kDa almost disappeared (Supplementary Figures 1d and e). The complex was purified by a two-step affinity chromatography (Strep-Tactin and Ni-NTA affinity chromatography) and size exclusion chromatography in presence of DDM (Figure 1b, Supplementary Figure 2a).

**Figure 1.**
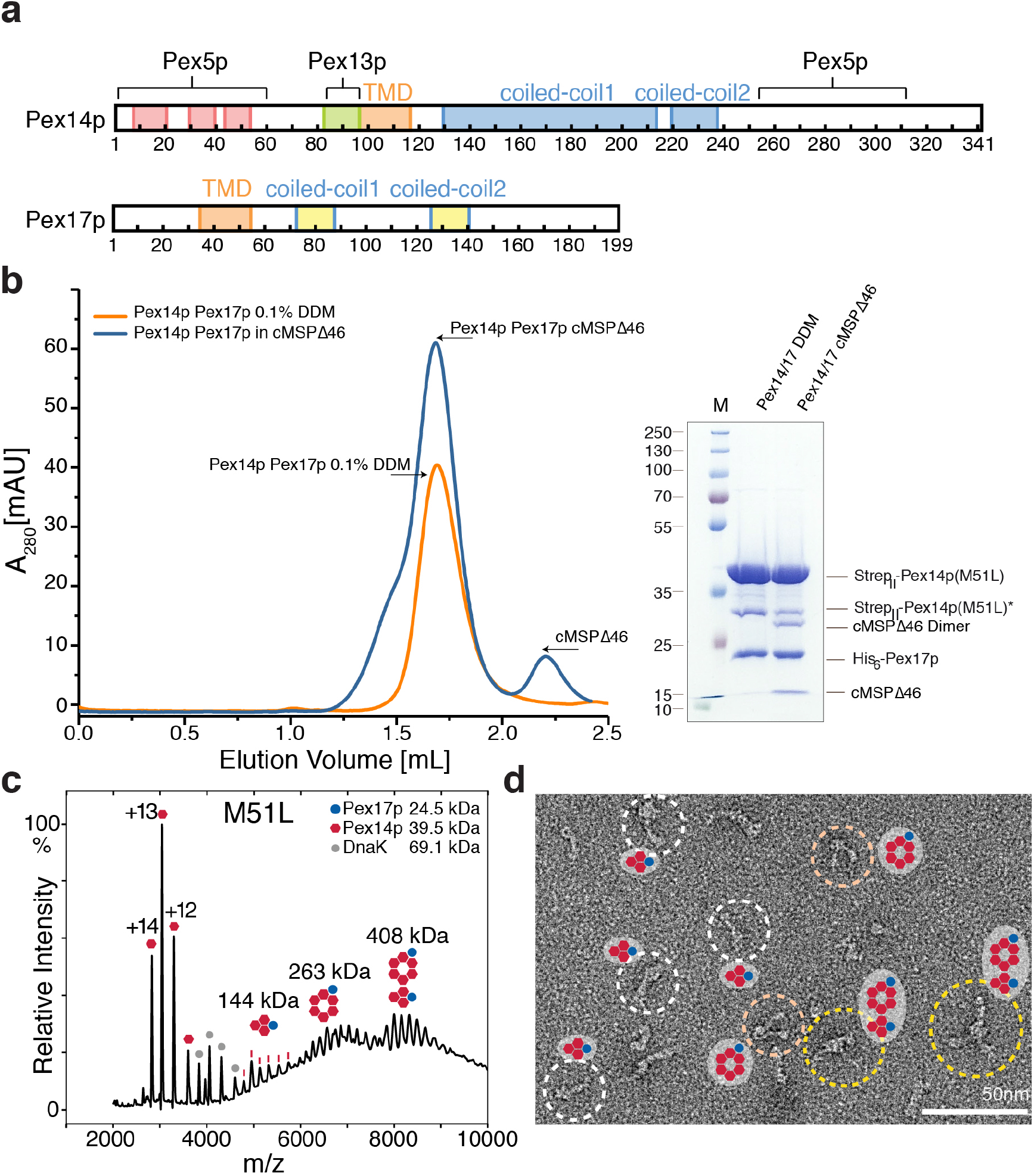
Molecular composition of Pex14p/Pex17p. **a)** Primary domain structure of yeast Pex14p and Pex17p. Red: alpha-helix; orange: predicted membrane domain, blue: predicted coiled-coil domain of Pex14p; yellow: predicted coiled-coil domain of Pex17p; pale green: Pex13p binding domain. Binding sites to the peroxisomal receptor Pex5p are also indicated. **b)** Size exclusion chromatography of recombinant Pex14p(M51L)/His_6_-Pex17p in DDM (orange) and cMSP1D1Δ4-6 nanodiscs (blue). Fractions of the Pex14p/Pex17p-peak used for subsequent EM studies are indicated by arrows and the respective SDS-PAGE is shown. **c)** Native MS spectra of recombinant Pex14p(M51L)/His_6_-Pex17p in DDM. Proteins were transferred into 200 mM ammonium acetate, pH 6.7 prior to native MS analysis. Peak series assigned to series of charge states are annotated by symbols as shown in the legend. Charge state series in the lower *m/z* region indicated masses of monomeric Pex14p and *E. coli* DnaK. Above *m/z* 4700 three non-resolved series were observed, best described by species of MW 144 kDa, 263 kDa and 408 kDa. Possible stoichiometries for the three masses are 3:1, 6:1 and 9:2 of Pex14p:Pex17p as schematically depicted above the spectrum. **d)** Representative subarea of a negative stain EM micrograph of recombinant Pex14p(M51L)/His_6_-Pex17p solved in an equivalent DDM concentration as used in MS measurements. White, orange and yellow dashed circles indicate monomers, dimers and trimers of the complex, respectively, explaining the different stoichiometries observed by native MS.

To investigate the composition of the complex, we then used native MS and gas phase collisional activation. Figure 1c shows native MS spectra for the coexpressed Pex14p(M51L)/Pex17p complex in DDM. The dominant peaks are assigned to a charge state series in the lower *m/z* region indicating masses of monomeric Pex14p and *E. coli* DnaK. Above *m/z* 4700, three partly resolved series were observed, which were assigned to molecular masses of 144 kDa, 263 kDa and 408 kDa. Isolation of ions within these series and subsequent collisional activation resulted in MS^2^ spectra showing charge state series indicative of masses of monomeric Pex14p and Pex17p (see Supplementary Figure 3). Possible stoichiometries to account for these three masses are therefore 3:1, 6:1 and 9:2 of Pex14p:Pex17p as schematically depicted the spectrum in Figure 1c. Of note, upon collisional activation, first monomeric Pex17p appeared as a main product, whereas at higher collisional energies Pex14p was predominant (Supplementary Figure 3).

Analysis of Pex14p(M51L)/Pex17p in DDM by negative stain EM revealed elongated flexible pin-like particles, which however displayed limited contrast (Supplementary Figure 2a). In accordance with the native MS data, the negative stain EM analysis revealed three distinct populations of particles: single rods, rod-dimers and rod-trimers (Figure 1d). The higher-order assemblies (rod-dimers and -trimers) can be explained by tail-to-tail hydrophobic interactions when the complex is solubilized in DDM.

We further used size-exclusion chromatography combined with multi-angle light scattering (SEC-MALS) to verify the molecular weight of Pex14p(M51L)/Pex17p. The complex indeed ran at 143 kDa (±0.02%) consistent with the 3:1 stoichiometry (Supplementary Figure 4). Based on these results, we conclude that the Pex14p(M51L)/Pex17p complex consists of three copies of Pex14p(M51L) and one copy of Pex17p, with a molecular weight of ~143-144 kDa.

We further isolated native complexes from oleic acid-induced cells expressing Pex14p-TPA, as previously described(*23*, *32*). Using Pex14p-TPA as bait, we isolated the native Pex14p/Pex17p complex from Triton X-100-solubilized whole cell membranes and analyzed the sample by negative stain EM (Supplementary Figure 5). This analysis confirmed that the native Pex14p/Pex17p complex features the same overall architecture as the recombinantly expressed complex (Supplementary Figure 5). Due to superior sample quality, further structural analyses were performed with recombinantly expressed protein complexes.

### CryoEM Structure of Pex14p(M51L)/Pex17p

To optimize the sample for EM studies and increase contrast, we replaced DDM with amphipol A8-35(*33*). However, the tight adsorption of amphipols to the hydrophobic transmembrane domains of Pex14p/Pex17p surprisingly resulted in dissociation of the complex and separation of Pex14p and Pex17p (Supplementary Figure 6). This might suggest that the transmembrane domains of Pex14p and Pex17p or regions in close proximity to them are involved in crucial interactions for Pex14p/Pex17p complex formation. To overcome this issue, we exploited the possibility to reconstitute the Pex14p/Pex17p complex into lipid nanodiscs. Both Pex14p and Pex17p contain a single transmembrane helix, thus depending on the oligomeric state of the complex, only few membrane-spanning helices are expected (Figure 1a). Therefore, we used the membrane scaffold protein (MSP)1D1dH5 with a rather narrow predicted scaffold diameter of 8-9 nm for nanodisc reconstitution. Despite extensive optimization efforts, the resulting proteo-nanodiscs were heterogeneous and incorporated multiple copies of the Pex14p/Pex17p complex, usually from both sides (Supplementary Figure 2b). We then repeated the reconstitution experiments using the recently reported circular MSP1D1Δ4-6 nanodiscs(*34*) (Figure 1b). The Δ4-6 circularized nanodisc with a diameter of 7 nm is according to our knowledge the smallest available nanodisc to date and, in addition, it shows superior stability and homogeneity in comparison to typical linear MSPs. Indeed, subsequent negative stain EM analysis revealed high-contrast monodispersed particles, with most nanodiscs trapping only a single copy of the complex that could be easily identified and automatically picked (Supplementary Figure 2c).

To select the structure of the Pex14p/Pex17p complex, we then recorded high-contrast images of Pex14p(M51L)/Pex17p reconstituted in MSP1D1Δ4-6 nanodiscs using Volta Phase-Plate (VPP) cryo-EM (Figures 2a and b). The digital micrographs show an even spread of elongated rod-like particles (Figures 2a and b), suggesting that most particles adopt a side-view orientation on the grid. Particles were selected automatically using crYOLO(*35*) and further processed with SPHIRE(*36*). Reference-free 2D class averages revealed a single copy of Pex14p/Pex17p within the nanodisc (Figure 2c). In contrast to other membrane proteins reconstituted in larger nanodiscs and analyzed by cryoEM(*37*, *38*), the protein in this case is not “floating” within the nanodisc, but rather inserted peripherally at close proximity to the MSP (Figure 2c). This might indicate possible interactions between the transmembrane domain of Pex14p and Pex17p and the scaffold helices. Transmembrane helices are however not resolved within the disc, suggesting either inaccurate alignment of the small transmembrane domain due to the compact density of the disc or flexibility of the transmembrane helices relative to the cytoplasmic rod. In contrast, the cytoplasmic domains are clearly visible and appear as five sequentially arranged globular densities and are stiff, mostly straight or slightly curved (Figure 2c). The class averages show an average length of ~220 Å (Figure 2d) and reveal additional flexibility at the end of the rod (Supplementary Video 1). We finally computed a 10 Å 3D reconstruction from ~82600 particles selected from the most homogeneous 2D class averages (Supplementary Figure 7).

**Figure 2.**
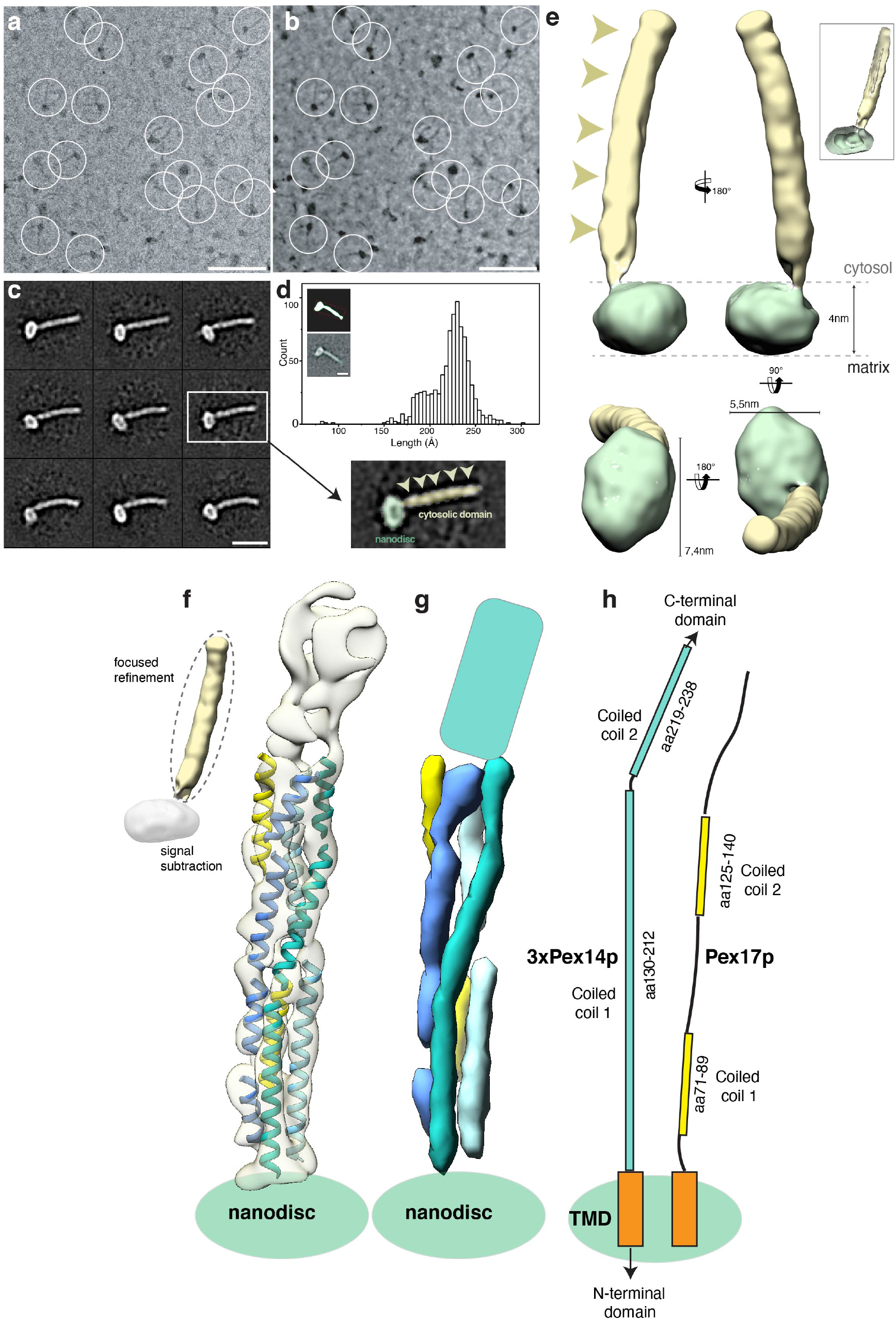
CryoEM structure of the Pex14p/Pex17p complex. **a)** Subarea of a typical low-dose VPP cryo-EM micrograph of Pex14p/Pex17p reconstituted in cMSP1D1Δ4-6. Some particles are highlighted with white circles. Scale bar, 50 nm. **b)** For better visualization we also denoised this particular micrograph using JANNI. It should be noted that denoised micrographs were not used during image processing. **c)** Representative reference-free 2D class averages of Pex14p/Pex17p. Scale bar, 10 nm. The inset shows one of the class averages further magnified. Densities of the nanodisc (green) and the cytosolic domains (yellow) are highlighted. **d)** End-to-end distance of the 2D class-averages. **e)** Density map of Pex14p/Pex17p in nanodisc. Shown are different side views, after horizontal rotation of 90° around their longest axis, the bottom and top view. The rod and nanodisc density, are highlighted in yellow and green, respectively. **f)** Resulting map from focused 3D local refinement on the rod region after signal subtraction of the nanodisc. The rod region dissembled in two structural regions, showing a kinked connection. poly-Ala helices are fitted into rod-like densities of the lower rod domain of the cryoEM map and colored according to their connectivity and assignment to Pex14p (shades of blue) and Pex17p (yellow) **g)** For better visualization and comparison, a density map was simulated from the poly-Ala-helices at a resolution of 8.5 Å. Fitting of α-helices was not possible in the upper rod domain (cyan), due to limited resolution. **h)** Schematic of the primary domain structures of yeast Pex14p and Pex17p showed according to their topology in their cryoEM structure (transmembrane domains: orange; coiled-coil domains Pex14p: cyan; coiled-coil domains Pex17p: yellow). The N- and C-terminal domains of Pex14p are not shown.

The final density map revealed a 22 nm elongated rod inserted at the periphery of the nanodisc (Figure 2e). The rod appears hollow with an inner diameter of 7 Å. We further performed focused refinement of the rod after signal subtraction of the nanodisc, which improved the respective density (Supplementary Figure 7a). The resulting structure can be dissembled in two structural regions, showing a kinked connection (Supplementary Figure 7b).

The resolution of the structural region below the kink is now significant, so that long α-helices are resolved, running approximately parallel to the rod axis (Figure 2f, Supplementary Video 2). In particular, three helices form a 120 Å coiled-coil-like bundle (Figures 2f-g). The helical bundle is however not well-ordered, showing several weak connections, suggesting thus a higher order degree of flexibility in comparison to a typical coiled coil structure. These weak connections do not involve however a change in the direction of the overall rod axis. Interestingly, two shorter α-helices complement (Figures 2f-g; yellow, Supplementary Video 2) this arrangement and integrate into the helical bundle. The overall length of 12 nm of each of the three long α-helices, as well as their topology directly after the membrane, agree well with the predicted coiled-coil domain 1 of Pex14p (Figures 2f-h, Figure 1a, Supplementary Figure 8). In addition, the dimensions and topology of the two shorter α-helices match the predicted coiled-coil domains 1 and 2 of Pex17p (Figures 2f-h, Figure 1a).

In accordance with the native MS and SEC MALS data suggesting a 3:1 heterotetrameric arrangement of Pex14p:Pex17p, we conclude that the lower part of the rod consists of a trimeric arrangement of the predicted coiled-coiled domain 1 of Pex14p (Pex14p_130-212_). Three copies of this domain are directly located after the membrane and form a 12 nm bundle, which is in addition complemented by a single copy of Pex17p, running parallel to the rod that is thus mainly formed by Pex14p. The two shorter coiled-coil domains of Pex17p (Pex17p_71-89_ and Pex17p_125-140_) are tightly associated with the bundle and the rod displays at the respective segments a pseudo-4-fold symmetry (Figures 2f-h, Supplementary Figure 7b, Supplementary Video 2).

The shorter structural region above the kink is not resolved, suggesting a higher degree of structural heterogeneity and/or flexibility (Figures 2f-h). However, the length of this part of the rod matches well the predicted coiled-coil domain 2 of Pex14p (Pex14p_219-238_) (Figures 2f-h)

### Interactions within *in vitro* and native Pex14p/Pex17p complexes

To obtain more detailed structural information on protein interactions within the Pex14p/Pex17p complex, the complex was subjected to chemical crosslinking combined with MS analysis (XL-MS). Figure 3a represents a schematic overview of the connected residues identified.

**Figure 3.**
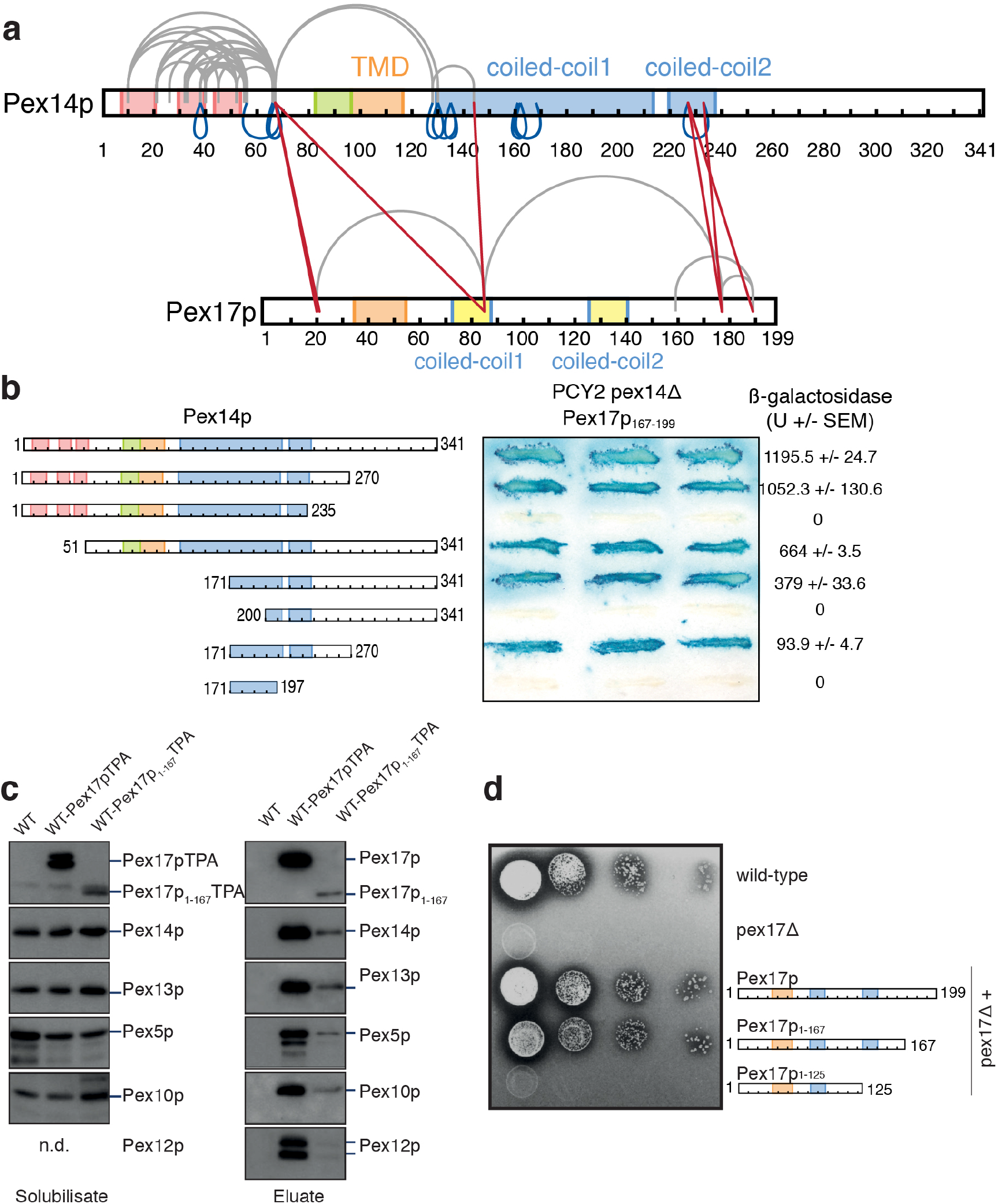
Characterization of the interactions between Pex14p and Pex17p. a) Map of connections between side chains of Pex14p and Pex17p identified by XL-MS. Purified Strep_II_-Pex14p/His_6_-Pex17p complexes were cross-linked using BS3 and subjected to in-solution digestion using trypsin. Straight red lines represent intermolecular crosslinks, grey curved lines intra molecular crosslinks and blue lines indicate homomultimeric linkages. **b)** Gal4p-binding domain (DB)/Pex14p fusions with progressive N- and/or C-terminal deletions of Pex14p were assayed for their interaction with Pex17p_167-199_. Double transformants of PCY2 *pex14*Δ cells expressing the indicated fusion proteins were selected, and ß-galactosidase activity was determined by a filter assay using X-gal as substrate. Three representative independent double transformants are shown. The ß-galactosidase activity shown on the right is the average of triplicate measurements for three independent transformants harboring each set of plasmids. SEM: Standard Error of Mean. U: Units **c)** Native complexes were isolated from solubilized membrane fractions from wild-type cells expressing either Pex17pTPA or Pex17p_1-167_TPA eluted from IgG-Sepharose with TEV protease. An eluate of wild-type cells served as control. Equal portion of the solubilisate (left) and eluate (right) were subjected to immunoblot analysis with antibodies as indicated. **d)** Cells of wild-type-, *pex17*Δ- or *pex17*Δ-*cells* expressing plasmid-based Pex17p-variants as indicated were grown overnight on glucose minimal media. Subsequently, 10-fold dilutions were prepared, and 2 μl of each dilution were spotted onto oleate plates, which were scored for the appearance of colonies and halo-formation. In contrast to wild-type cells, *pex17*Δ mutant was not able to utilize oleate. Expression of wild-type Pex17p as well as Pex17p_1-167_ restored the growth defect of *pex17*Δ cells, whereas Pex17p_1-125_ lost its ability for functional complementation.

Blue lines in Figure 3a represent identified homomultimeric linkages that are crosslinks involving two overlapping peptides, and thus, originating from two molecules of Pex14p. Gray lines in Figure 3a represent intramolecular contacts and most of them span the N-terminal domain of Pex14p. This is in agreement with the previous crystal structure of the conserved N-terminal domain of Pex14p with a three-helix bundle(*39*). Pex14p-Pex14p linkages (blue lines; Figure 3a) span the full region of predicted coiled-coils and are further localized within and close to the N-terminal three-helix domain of Pex14p, indicating a parallel arrangement between adjacent Pex14p molecules. Interestingly,Pex14p and Pex17p display intermolecular linkages (red lines; Figure 3a) along their complete sequence.

These data are in agreement with the cryoEM structure, indicating not only a parallel arrangement between adjacent predicted coiled-coil domains of Pex14p, forming thereby the helical extramembrane coiled-coil rod, but also a parallel arrangement and interactions between adjacent coiled coil domains of Pex14p and Pex17p molecules.

Several linkages indicate major contacts between the C-terminal end of Pex17p (aa177-189) and the second predicted coiled-coil of Pex14p. This suggests that the rod-like domain above the kink, that is not as well resolved in our cryoEM structure, is apparently formed by three copies of coiled-coil domain 2 of Pex14 (Figures 2f-h) and contains in addition the unstructured C-terminal end of Pex17p, most likely in a parallel arrangement relative to Pex14p. These linkages are in agreement with previous results identifying the extreme C-terminus of Pex17p (corresponding to aa167-199) as the smallest fragment that is sufficient for Pex14p-interaction(*40*). Additional interfaces were identified between the N-terminal domains of Pex14p (Lys67) and Pex17p (aa20-21), indicating further secondary contacts between both proteins.

To further characterize the Pex14p-Pex17p interactions, we performed domain-mapping experiments based on the yeast two-hybrid system with Pex17p_167-199_ fused to the Gal4p-DNA-binding domain (DB) and Gal4p-DNA-activation domain fusions of progressive carboxyl- and/or amino-terminal truncations of Pex14p. In line with previous findings, co-expression of Pex17p_167-199_ with fulllength Pex14p resulted in considerable ß-galactosidase, demonstrating the interaction of both proteins(*40*) (Figure 3b). However, Pex14p lacking N- or C-terminal regions did still interact with Pex17p_167-199_, pointing to an internal binding region. In fact, we identified the second putative coiled-coil domain of Pex14p plus a C-terminal portion as sufficient to mediate interaction to Pex17p_167-199_ (Figure 3b). This is in line with the XL-MS data, which also revealed the second putative coiled-coil domain of Pex14p as a major interaction site with Pex17p (red lines, Figure 3a).

Of note, Pex17p lacking its C-terminus did not interact with Pex14p in yeast two-hybrid assays(*40*), although further N-terminal contacts were detected by XL-MS (Figure 3a). Thus, in order to characterize possible secondary binding sites, Pex17p lacking its C-terminus (Pex17p_1-167_) was analyzed in its endogenous environment, the peroxisomal membrane, instead of the nucleus, as it is the case in yeast two-hybrid assays. To this end, a wild-type strain expressing either Pex17p or Pex17p_1-167_ genomically tagged with protein A (Pex17pTPA, Pex17p_1-167_TPA) with a TEV cleavage site between both fusion partners was used. In line with published data and the function of Pex17p as a constituent of the receptor docking complex, Pex13p, Pex14p as well as the PTS1-receptor Pex5p were co-isolated when full-length Pex17p served as bait (Figure 3c)(*41*). As originally described(*23*), other components of the importomer, represented by the two RING-finger proteins Pex10p and Pex12p, were also part of the Pex17p-complex. Pex17p lacking its C-terminal Pex14p-binding module displayed a significant reduction in its steady-state level when compared with wild-type protein and the constituents of the importomer that were present in the Pex17p-complex, were also associated with Pex17p_1-167_TPA, albeit in a much lower amount (Figure 3c). Thus, the results demonstrate that Pex17p_1-167_ is still associated with the importomer, although it is lacking its C-terminal Pex14p-binding region. This is again in line with the XL-MS results, corroborating that interactions between the first putative coiled-coils and between the N-termini of co-expressed Pex14p and Pex17p (Figure 3a) promote the insertion of Pex17p_1-167_ into Pex14p membrane complexes (Figure 3c). Interestingly, growth on oleic acid of cells deficient in Pex17p was restored upon expression of Pex17p_1-167_, indicating that the truncated Pex17p_1-167_ is biologically active (Figure 3d).

### Intermolecular interactions within the predicted coiled-coil domains of Pex14p are essential for its oligomerization

To assess which of the Pex14p-Pex14p interactions are essential for oligomerization, we used recombinantly expressed truncated Pex14p-variants and monitored formation of oligomers by chemical crosslinking and native MS. One N-terminal variant, Pex14p_1-95_, and two C-terminal variants comprising solely the second coiled-coil domain (Pex14p_213-341_) or both coiled-coil domains (Pex14p_119-341_) were incubated with the crosslinker BS3 and products of the reactions separated by SDS-PAGE (Figure 4a). Whereas the N-terminal and the shorter C-terminal variant remained in the monomeric form, the larger C-terminal variant including both coiled-coil domains showed additional bands indicative of the formation of dimers and trimers (Figure 4a). With increasing concentrations of the crosslinker, the trimer band became stronger than the dimer band, but no higher molecular weight bands of tetrameric or higher oligomeric species appeared (Supplementary Figure 9). The same Pex14p variants were studied by native MS preserving non-covalent interactions during transfer into the gas phase and subsequent analysis (Figure 4b). In line with the crosslinking results, the two shorter variants were observed solely monomeric, whereas the longer variant comprising both coiled-coil domains predominantly formed trimers and, to a considerably lesser extent, dimers and monomers.

**Figure 4.**
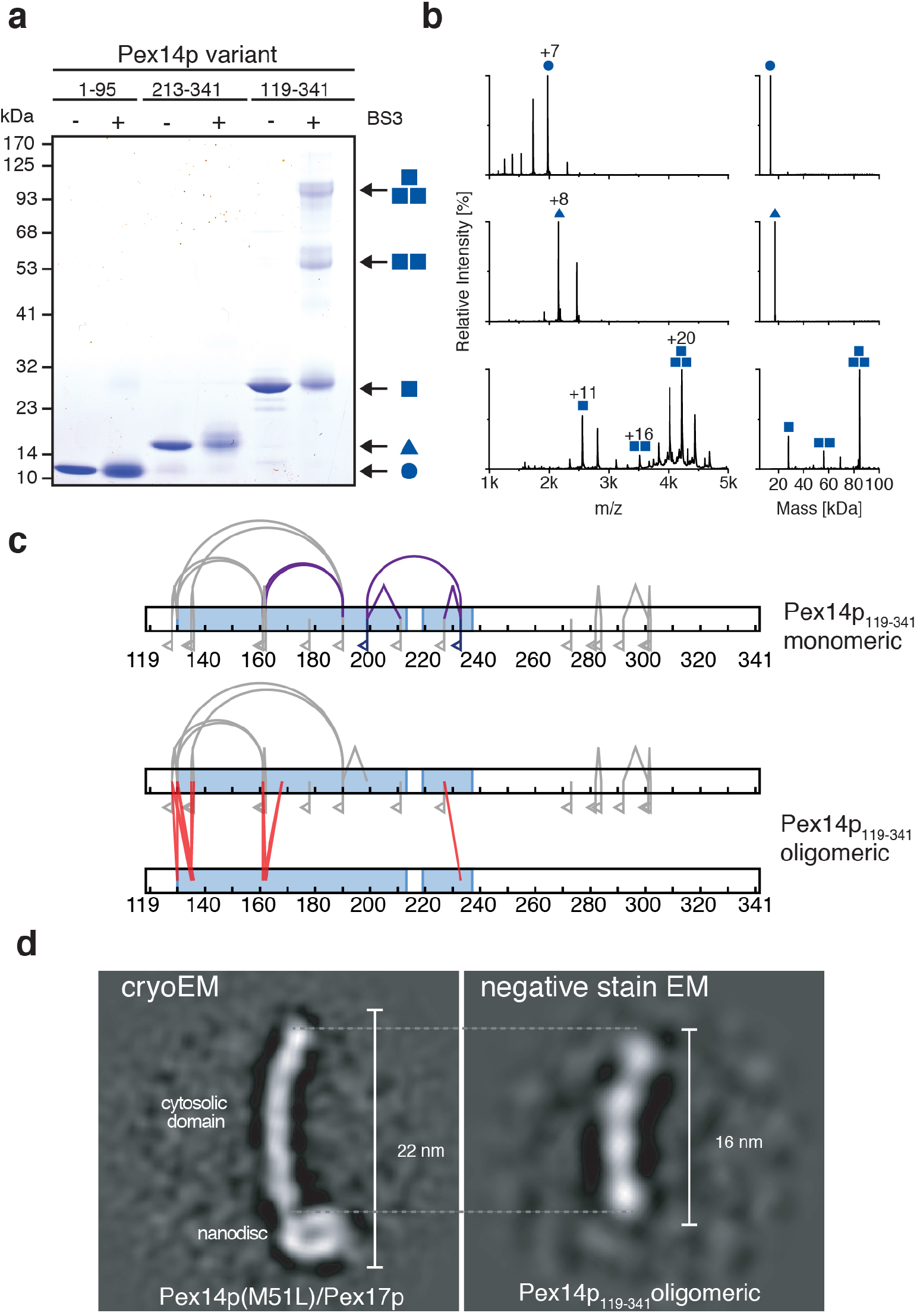
Oligomerization properties of Pex14p variants. **a)** Recombinant variants of Pex14p comprising its N-terminal domain (Pex14p_1-95_; circle), a shorter (Pex14p_213-341_; triangle) and a longer C-terminal part (Pex14p_119-341_; square) were cross-linked using BS3 and separated by SDS-PAGE. Bands assigned to monomeric, dimeric and trimeric forms are marked by one, two or three symbols. **b)** Native MS of Pex14p_1-95_ (top), Pex14p_213-341_ (middle) and Pex14p_119-341_ (bottom spectrum). Proteins were transferred into 200 mM ammonium acetate, pH 6.7, and analyzed by nano-ESI-QTOF-MS. Main charge states are indicated based on assignments of peaks series to monomeric and oligomeric complexes as indicated. The respective charge-state deconvoluted spectra are shown on the right. **c)** Schematic view of intermolecular (straight red lines) and intramolecular (grey curved lines) crosslinks as well as peptide modifications by loop-links (grey angled lines) and mono-links (flags) identified and quantified by LC-MS in monomeric (top) and oligomeric (bottom) Pex14p_119-341_. Quantitative analysis further distinguished linkages recovered equally from monomeric and oligomeric forms (grey) and monomer-specific linkages (purple). **d)** Side-by-side comparison between reference-free class averages of Pex14p(M51L)/His_6_-Pex17p in nanodisc (cryoEM) and the homotrimeric Pex14p_119-341_ variant (negative stain).

From the crosslinking experiment, gel-separated monomeric and oligomeric forms of Pex14p_119-341_ were subjected to digestion using trypsin and LC-MS/MS analysis for peptide identification. In addition, a quantitative MS analysis was performed, enabling a direct comparison of the abundance of individual crosslinked peptides identified in the homo-oligomer to those in the monomer (Supplementary Figure 10). The map of interactions resulting from this analysis is summarized in Figure 4c. We identified a set of 13 residue pairs specifically enriched with the homo-oligomer representing intra-molecular interactions along the full region of both predicted coiled-coil domains (Supplementary Figure 10c). Only four residue pairs had ratios close to 1:1 and, thus, were present in almost equal abundance in monomers and oligomers. Of note, one residue pair of high abundance (199/233) and two pairs of lower abundance (162/190 and 161/190) were monomer-specific, suggesting that these linkages may prevent assembly into oligomeric forms.

In conclusion, *in vitro* formation of Pex14p oligomers (predominantly homotrimers) required the presence of its first coiled-coil domain. The identified intermolecular interactions along the full length of both predicted coiled-coil domains of Pex14p homo-trimers confirm the parallel assembly of the individual molecules. Connections between the two coiled-coil domains within Pex14p monomers in absence of the transmembrane domain interfere with the formation of higher oligomers, possibly by forcing bended conformations not compatible with the parallel geometry.

We further subjected the longer trimeric variant (Pex14p_119-341_ homotrimers including the two coiled-coil domains and the C-terminal domain of Pex14p) to negative stain analysis. The trimeric subcomplex forms 16 nm rods that appear approximately like the full-length Pex14p/Pex17p complex without the nanodisc density (Figure 4d, Supplementary Figure 11). This further confirms that three parallel arranged predicted coiled-coil domains of Pex14p indeed assemble to form the backbone of the elongated extra-membrane rod-like domain of the Pex14p/Pex17p complex.

### Membrane topology of Pex14p/Pex17p

At this point, it is important to emphasize that yeast Pex14p contains two binding sites for the PTS1 receptor Pex5p: the first at the N-terminal domain and the second at the C-terminal domain (Figure 1a). Both domains are not resolved in our cryoEM structure. The C-terminal domain of Pex14p (~100 aa) is predicted to be intrinsically disordered and, thus, highly flexible, whereas the N-terminal domain is highly conserved and forms a three-helix bundle(*39*), which is however connected to the transmembrane domain by a long flexible linker peptide (Supplementary Figure 8), allowing apparently a high degree of flexibility. With regard to Pex17p, only the helices forming the two-short coiled-coil domains above the membrane are identified in our cryoEM structure.

To verify the presence of the N-terminal domains of both Pex14p and Pex17p within the rod complexes observed by cryoEM and clarify their membrane topology, we co-expressed (Pex14p/His_6_-Pex17p) and (His_6_-Pex14p/Pex17p),reconstituted the respective complex in MSP1D1Δ4-6 nanodiscs, labelled with Ni-NTA-gold and imaged by negative stain EM (Figures 5a and b). In both cases, the gold particles bound below the nanodiscs, suggesting co-localization of the N-termini of Pex14p and Pex17p. This is in agreement with the XL-MS data that revealed linkages between the N-terminal domains of Pex14p (Lys67) and Pex17p (aa20-21) (Figure 3a). Our data clearly suggest that the two binding sites of Pex14p for the cargo-loaded receptor are apparently located on opposite sides of the membrane. The first N-terminal site is located below the nanodisc, whereas the second C-terminal site is located after the extramembrane rod, above the nanodisc.

**Figure 5.**
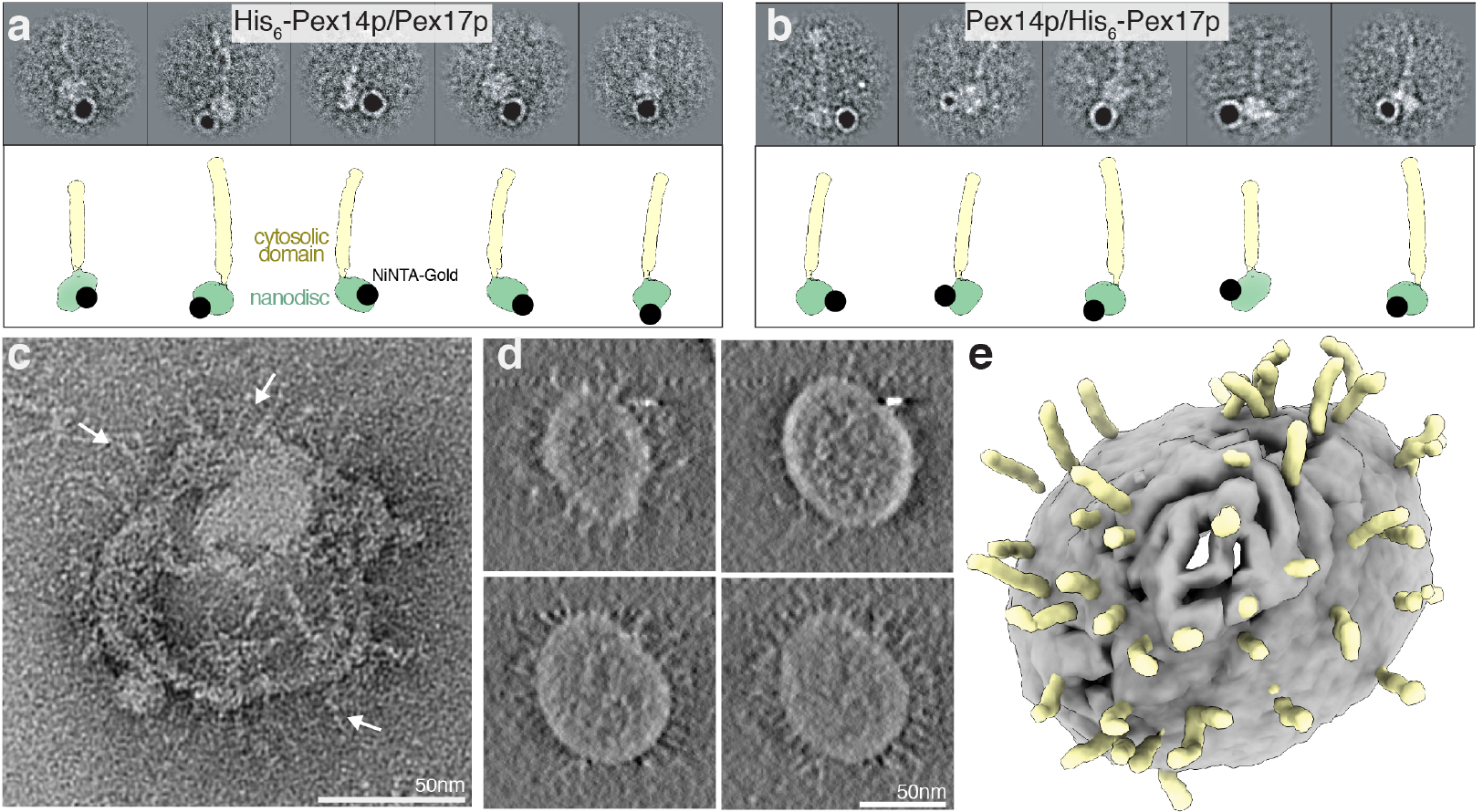
Immunogold labeling of the N-termini of Pex14p/Pex17p and incorporation into liposomes. **a-b)** Localization of the N-terminal His_6_-tag in the recombinant His_6_-Pex14p(M51L)/Pex17p (a) and Pex14p(M51L)/His_6_-Pex17p (b) with Ni-NTA-nano-gold. The topology of the gold label is also shown graphically mapped on the model of Pex14p/Pex17p. The N-termini of both proteins are clearly located closely to the nanodisc. **c)** Representative negative stain image of Pex14p/Pex17p incorporated into liposomes. **d-e**) Slices through the tomographic cryo-EM volume of a single Pex14p/Pex17p-liposome (d) and corresponding 3D segmentation (e), showing the bilayer (gray) and the characteristic Pex14p/Pex17p rods (yellow).

In order to exclude the possibility that the narrow nanodiscs prevent the formation of higher order assemblies and challenge the oligomerization and overall structure of the Pex14p/Pex17p complex in a more close-to-native lipid environment and membrane curvature, we reconstituted the complex in liposomes and visualized the liposomes by negative stain EM and cryoET (Figures 5c-e). After successful reconstitution, the liposomes were shaped like sea urchins, with multiple copies of rods (spikes) covering isotropically their outer surface. 20 nm single rods were visible around the circumference of the liposomes mostly arranged perpendicular to the bilayer plane, clearly resembling the cryo-EM structure in the nanodisc (Figures 5c-e, Supplementary Video 3). Pex14p/Pex17p do not form larger super-complexes upon liposome incorporation. Taking into account recent studies indicating an intra-peroxisomal localization of the N-terminal domain of Pex14p(*30*), we also conclude that the Pex14p/Pex17p rods (Figure 5e) are facing the cytosol.

## Discussion

It is well established that Pex13p, Pex14p, and Pex17p mediate the docking of cytosolic receptor-cargo complexes to the peroxisomal membrane(*10*–*14*, *42*–*44*). Pex13p and Pex14p recognize and physically bind both of the import receptors, Pex5p and Pex7p. This, and the fact that Pex17p interacts with Pex14p(*14*, *44*), reviewed in (*45*) led to the conclusion that these three peroxins form the receptor-docking complex. However, docking of the receptor cargo complex supposedly is mediated by Pex14p(*21*, *22*). Along this line, transport of the peroxin Pex8p into peroxisomes, requires only the presence of Pex14p and the import receptor Pex5p(*46*). Moreover, *in vitro*, Pex5p and Pex14p alone are capable to form a gated ion-conducting channel(*15*). Taken together, these data suggest that Pex14p oligomers constitute minimal receptor-docking complexes. In our study, we characterize the overall architecture of Pex14p in complex with Pex17p. Pex17p is present only in yeast and in this case, coexpression of Pex14p with Pex17p in *E. coli* was crucial to obtain samples of sufficient quality and amount for structural studies.

Our study provides the first insights into the architecture of this central component of the peroxisomal import machinery in a membrane-mimicking environment, which we summarize in Figure 6a. We show that Pex14p and Pex17p together form a ~140 kDa complex (Figure 1c), containing three copies of Pex14p and one copy of Pex17p. The complex resembles a rod-like particle with one end inserted in the lipid nanodisc (Figures 2c and e). The overall length of the rod-part (~18 nm) corresponds approximately to the predicted length of the two elongated Pex14p coiled-coil domains. We attempted to solve the structure of the complex by single particle cryo-EM, but visualization of a thin helical bundle inserted into, according to our knowledge, the smallest available lipid nanodisc, was not possible for conventional defocus-based imaging. Therefore, we used the Volta Phase Plate(*47*) to enhance the contrast of this weak phase object. We were able to obtain a low resolution reconstruction of the complex (Figure 2e) and further focused refinement of the rod after signal subtraction allowed us to reach sub-nanometer resolution, revealing the helical arrangement of the lower structural region of the rod (Figure 2f; Supplementary Video 2). These data are in line with the 3:1 stoichiometry of the complex, suggesting a homo-trimeric coiled-coil core formed by the predicted coiled-coil domain 1 of three copies of Pex14p and further association of both predicted coiled-coil domains of a single Pex17p copy into this arrangement (Figure 6a). At the respective segments, the rod shows however several kinks and does not display a typical tetrameric coiled-coil structure, as we initially expected, allowing thereby possibly a higher degree of flexibility.

**Figure 6.**
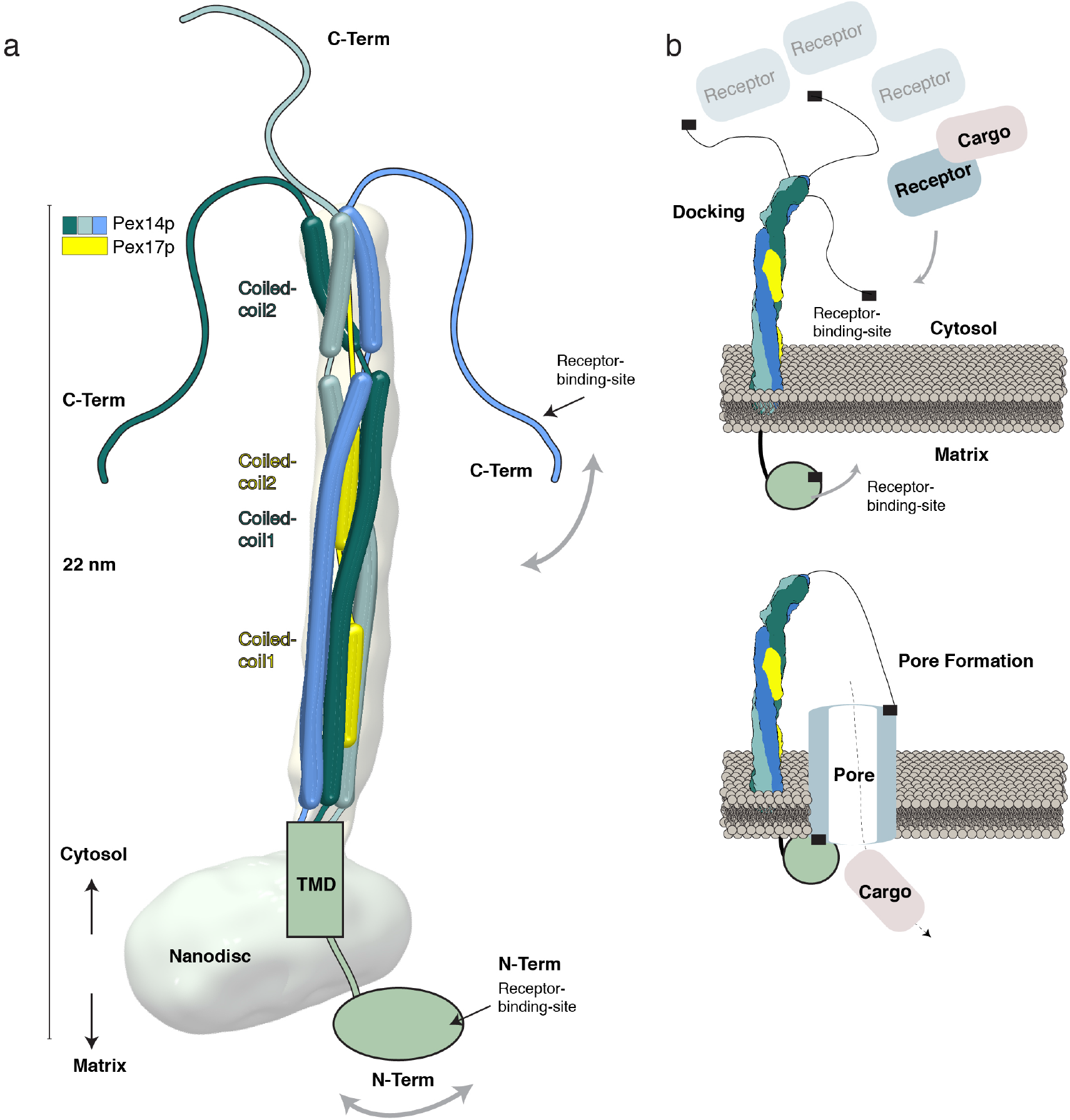
Organization of the Pex14p/Pex17p complex. **a)** Model of the Pex14p/Pex17p complex. The topology and arrangement of the different domains are indicated, according to the results of the present study. **b)** Model for triggering peroxisomal translocation upon binding of the receptor-cargo complex to Pex14p/Pex17p at the peroxisomal membrane.

The XL-MS data identified intermolecular interactions along the full length of Pex14p and Pex17p (Figure 3a). Mapping of the Pex14p-Pex17p interactions was further confirmed by yeast two-hybrid assays and functional expression of a Pex17p variant (Figures 3b and c). The data clearly suggest a parallel arrangement of both molecules along their complete sequence (Figure 6a). Both coiled-coil domains of Pex14p are required for homooligomerization (Figure 4). Importantly, the three copies of the coiled-coil domains of Pex14p show a parallel and not an anti-parallel arrangement relative to each other. This trimeric parallel coiled-coil arrangement of Pex14p_119-341_ forms indeed the backbone of the extra-membrane rod of the Pex14p/Pex17p complex (Figure 4d).

The transmembrane domains of Pex14p and Pex17p could not be resolved, even after signal subtraction and focused refinement on the nanodisc density, suggesting a flexible arrangement of four α-helices spanning the membrane. The conserved N-terminal domain of Pex14p is connected to the transmembrane domain by a flexible linker (Supplementary Figure 8, Figure 6a). Indeed, two-dimensional class averages of Pex14p/Pex17p indicate that this part of the complex is highly flexible, only appearing as blurry density below the nanodisc (Supplementary Video 1). Nevertheless, gold labeling experiments allowed us to detect the N-termini of both Pex14p and Pex17p below the nanodisc, suggesting an intra-peroxisomal localization of the respective domains (Figure 5, Figure 6a). This is in agreement with recent studies, suggesting a N_in_-C_out_ topology of Pex14p(*30*). CryoEM tomograms of reconstituted Pex14p/Pex17p in liposomes revealed isotropic decoration of the liposomes by the complex resulting in a sea urchin-like proteo-liposome. Thus, the Pex14p/17p complexes do not locally modify membrane curvature and do not form higher-order assembly structures or clusters in a more close-to-native lipid environment and membrane curvature.

According to our results, the two receptor binding sites of Pex14p are located on opposite sides of the peroxisomal membrane: the first at the N-terminal domain (intra-peroxisomal) and the second at the C-terminal domain (facing the cytosol) (Figure 6a). The C-terminal domain of Pex14p (~100 aa) is not resolved in our cryo-EM structure. However, according to secondary and 3D structure predictions (Supplementary Figure 8), this domain containing the cytosolic receptor-binding-domain is highly disordered and apparently flexible (Figure 6a). Importantly, according to our XL-MS analysis, the C-terminal domains of Pex14p do not show homomultimeric-linkages (Figure 3a; Figure 6a).

Our data thus strongly indicate that the peroxisomal docking complex, mainly formed by Pex14p, resembles a rod-like structure that emanates into the cytosol and uses highly flexible terminal peptides to recruit the cargo-loaded receptor complexes (Figure 6b). The further steps of pore formation and cargo translocation can only be speculated at this stage. The overall Pex14p architecture presented in this study renders the possibility of dramatic conformational changes of the Pex14p coiled-coil core and entry of further Pex14p domains into the membrane upon receptor binding rather unlikely. In particular, the 20 nm long and compact assembly of its central coiled-coil domain is not consistent with dramatic structural rearrangements of this domain. A cluster of charged residues in-between the transmembrane helix and the first coiled-coil helix (Supplementary Figure 8) would prevent insertion of the coiled-coil helices into the membrane bilayer. The possible distance of the C-terminal receptor binding site to the membrane is unlikely to allow entry of this unstructured domain into the membrane without dramatic conformational changes of the coiled-coil domain. Furthermore, the α-helices of the conserved intra-peroxisomal flexible N-terminal domain containing the first receptor binding site are not amphipathic, and thus not capable to enter the membrane (Figure 6b). The overall linear antenna-like architecture of the complex suggests rather a pivotal role of Pex14p in recruitment of Pex5p/cargo complexes and providing the structural support necessary for further formation of the translocation pore. This would imply that the pore may be exclusively formed by the receptor Pex5p, which is in line with the transient pore model(*48*) suggesting that Pex5p is functioning like a pore-forming toxin (Figure 6b). Indeed, import-deficient *pex14*Δ yeast mutants are capable to import matrix proteins after Pex5p over-expression(*49*) and *in vitro*, Pex5p is capable to enter the membrane without any help of the docking complex, when present at high concentrations(*50*). The interaction of the N-terminal domain of Pex14p to membrane-embedded Pex5p might then further stabilize the Pex5p transient pore assembly (Figure 6b).

Our results provide the first mechanistic insight into the yeast peroxisomal docking complex and thus into major aspects of the peroxisomal import machinery in general. Achieving near atomic resolution is beyond our grasp now, but we are confident that our study will provide a strong foundation and trigger future *in vivo* and *in vitro* structural and integrative studies of higher order assemblies, towards understanding key events of peroxisomal translocation.

## Acknowledgments

We are grateful to D. Prumbaum for excellent assistance with electron microscope facilities and S. Tacke for acquisition of the tomographic tilt series. We thank Raphael-Gasper-Schoenenbruecher for his excellent support during SEC-MALS data collection and analysis, as well as fruitful discussions. We also thank Sven Fischer for experiments during the initial phase of this work, Bettina Knapp for technical assistance with LC/MS analyses as well as Carla Schmidt and Jan Commandeur (MS Vision) for discussions on native MS. C.G thanks Stefan Raunser for continuous support and I. Vetter for stimulating discussions. This work was supported by the Max Planck Society, the Deutsche Forschungsgemeinschaft (DFG, German Research Foundation) (FOR1905 to C.G., R.E, B.W). Work in the Warscheid lab was also funded by the DFG – Project-ID 278002225 – RTG 2202 and – Project-ID 403222702 – SFB 1381, the Excellence Strategy (CIBSS – EXC-2189 – Project-ID 390939984) and the Excellence Initiative of the German Federal & State Governments (EXC 294, BIOSS). Work included in this study has also been performed in partial fulfillment of the doctoral theses of P.L., T.H. and D.W..

## Accession codes

The EM density map has been deposited in the Electron Microscopy Data Bank under accession code EMD-XXX. MS raw data and result files have been deposited to the ProteomeXchange Consortium via the PRIDE repository(*51*) and are publicly accessible from its website (http://www.ebi.ac.uk/pride) with the dataset identifier PXD016304.

## Author contributions

T.H. and W.G. designed proteins, performed mutational studies and yeast-two hybrid assays. P.L., T.H., D.W., T.J. purified proteins; P.L., T.H. optimized proteins for structural studies; J.M. assisted protein reconstitution in circular nanodics; P.L., B.K. screened samples and collected negative stain data; P.L. collected cryo electron microscopy data; P.L. and C.G. processed, refined and analyzed cryo-EM SPA and cryoET data; D.W. optimized proteins for MS studies; D.W., J.B. and F.D. performed crosslinking MS and native MS experiments and analyzed MS data; C.G. wrote the manuscript; R.E., B.W. co-wrote the manuscript. C.G., R.E., B.W. designed the study and managed the project. All authors discussed the results and commented on the manuscript.

---

**Supplementary Figure 1.**
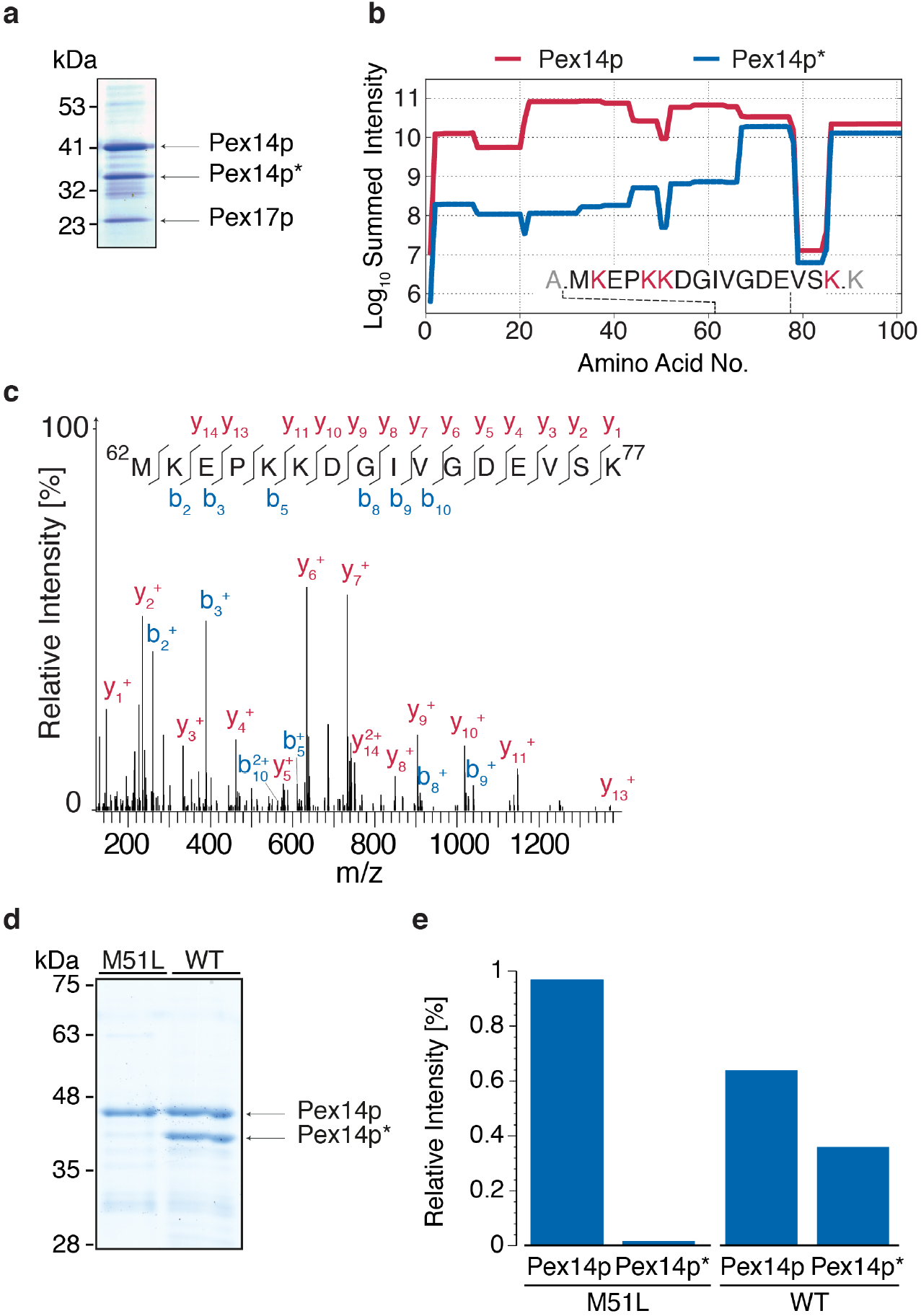
Partial degradation of recombinant Pex14p by cleavage at methionine 51 was prevented by the mutation M51L. **(a)** Strep_II_-Pex14p/His_6_-Pex17p-complex analyzed by SDS-PAGE revealed a dominant Pex14p-degradation product (Pex14p*) at an apparent molecular mass of 35 kDa. **(b)** Intensities measured by LC-MS of tryptic peptides recovered from individual gel bands of Pex14p (red line) and Pex14p* (blue line) indicate that the N-terminal part up to methionine 51 (M62 in Strep_II_-Pex14p) was absent in the degradation product. The inset shows the identified peptide representing the putative N-terminus of Pex14p*. Predicted cleavage sites of trypsin are highlighted in red. **(c)** MS/MS spectrum identifying the peptide generated by cleavage N-terminal to methionine 51. Fragment ions of type y (red) and type b (blue) identifying the peptide sequence are marked. **(d)** Recombinant Pex14p, wildtype and M51L were analyzed by SDS-PAGE and Coomassie staining. Proteins in individual gel slices were subjected to in-gel digestion using trypsin and quantitative LC-MS/MS analysis. The band assigned to the degradation product (Pex14p*) was significantly decreased in the M51L compared to wild-type. **(e)** LC-MS intensities of Pex14p peptides recovered from Pex14p* gel slices amounted to 1.6% and 36.1% of the total intensity for both gel slices for M51L and wild-type, respectively.

**Supplementary Figure 2.**
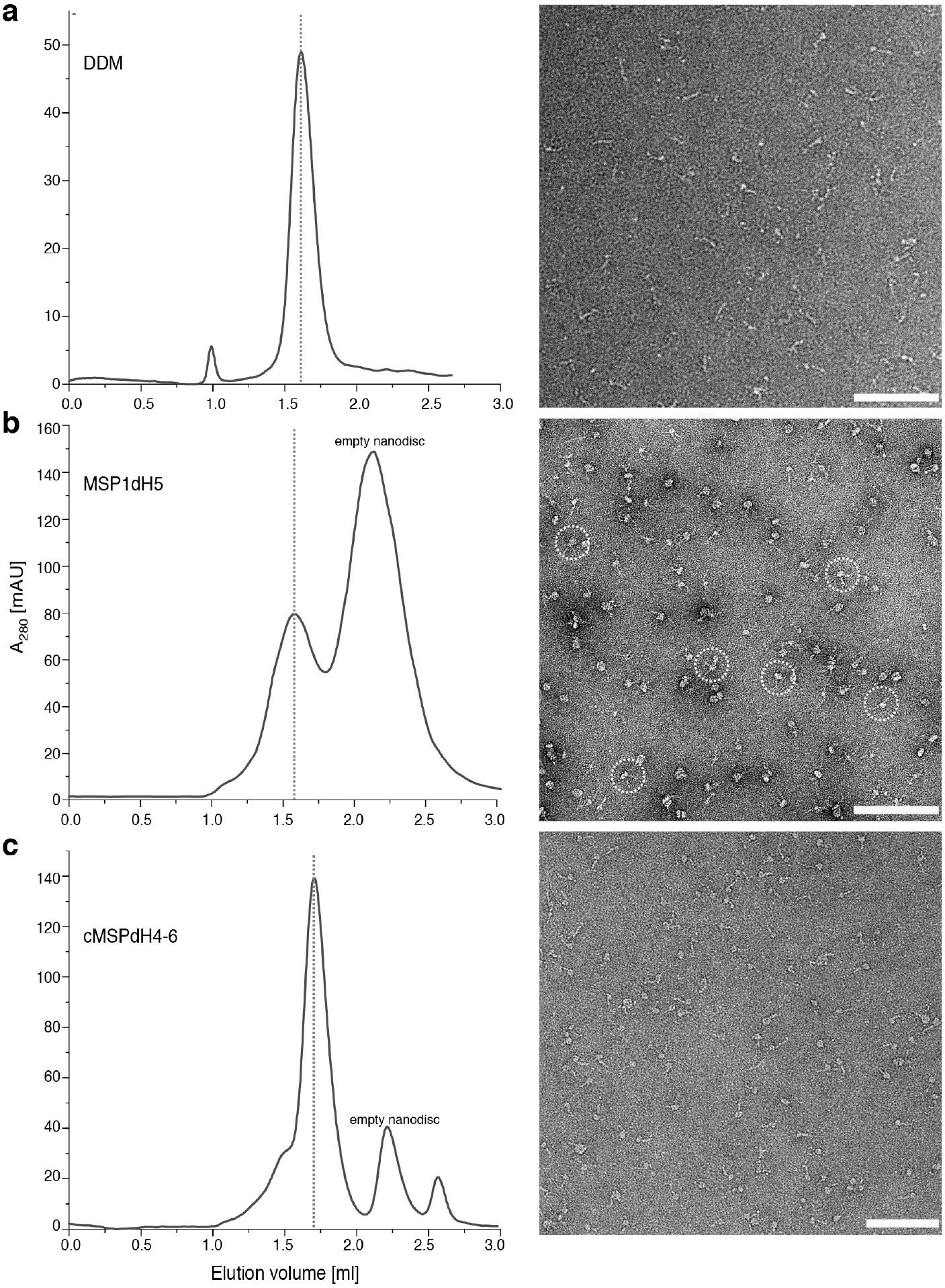
Size-exclusion chromatography and negative stain EM analysis of Pex14p/Pex17p. Gel-filtration chromatogram (left panel) and typical EM micrograph (right panel) of Pex14p/Pex17p in 0.1 % DDM **(a),** (MSP)1D1dH5 nanodiscs (ratio 1:30) **(b)** and MSP1D1Δ4-6 circular nanodiscs (ratio 1:30) **(c)**. Dashed lines indicate fractions used for EM. Dashed circles in (b) indicate proteo-nanodiscs including multiple copies of Pex14p/Pex17p. Scale bars 100 nm.

**Supplementary Figure 3.**
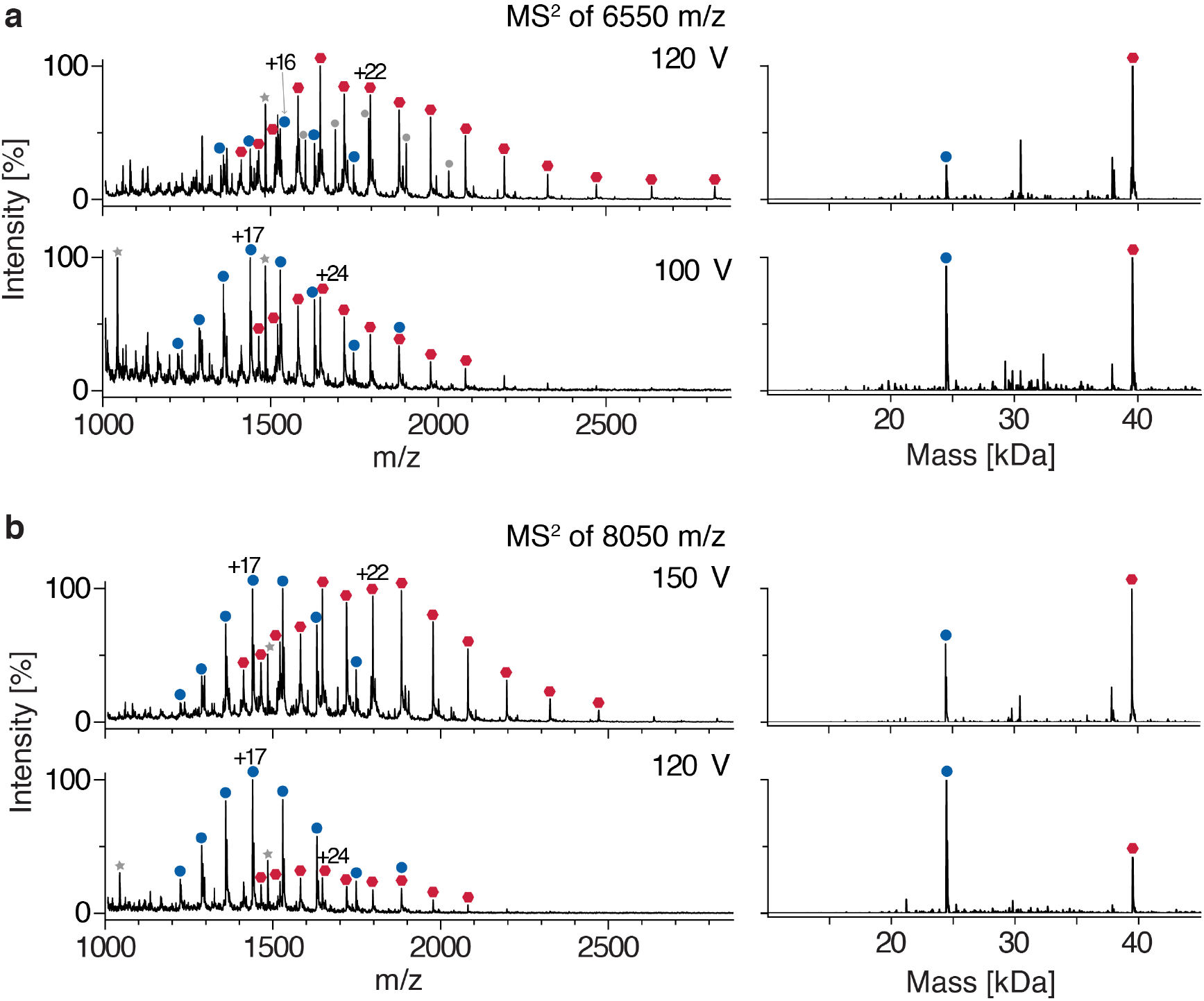
Native MS analysis of recombinant Strep_II_-Pex14p(M51L)/His_6_-Pex17p. **(a)** MS/MS spectra for the Strep_II_-Pex14p(M51L)/His_6_-Pex17p complex, generated by selection of ions of *m/z* 6550 in the quadrupole and subsequent collisional activation by acceleration voltages of 100 V (bottom) or 120 V (top). At a lower acceleration voltage of 100 V, the series corresponding to the mass of Pex17p (blue circles) is predominant, whereas at higher energy (120 V) Pex14p became more abundant (red hexagons). The right part shows the corresponding charge-state deconvoluted spectra. **(b)** Equivalent to (a), but after selection of ions of *m/z* 8050 in the quadrupole. Similarly, at a lower collision energy (120 V), Pex17p was observed as the predominant product, whereas at 150 V Pex14p was more abundant.

**Supplementary Figure 4.**
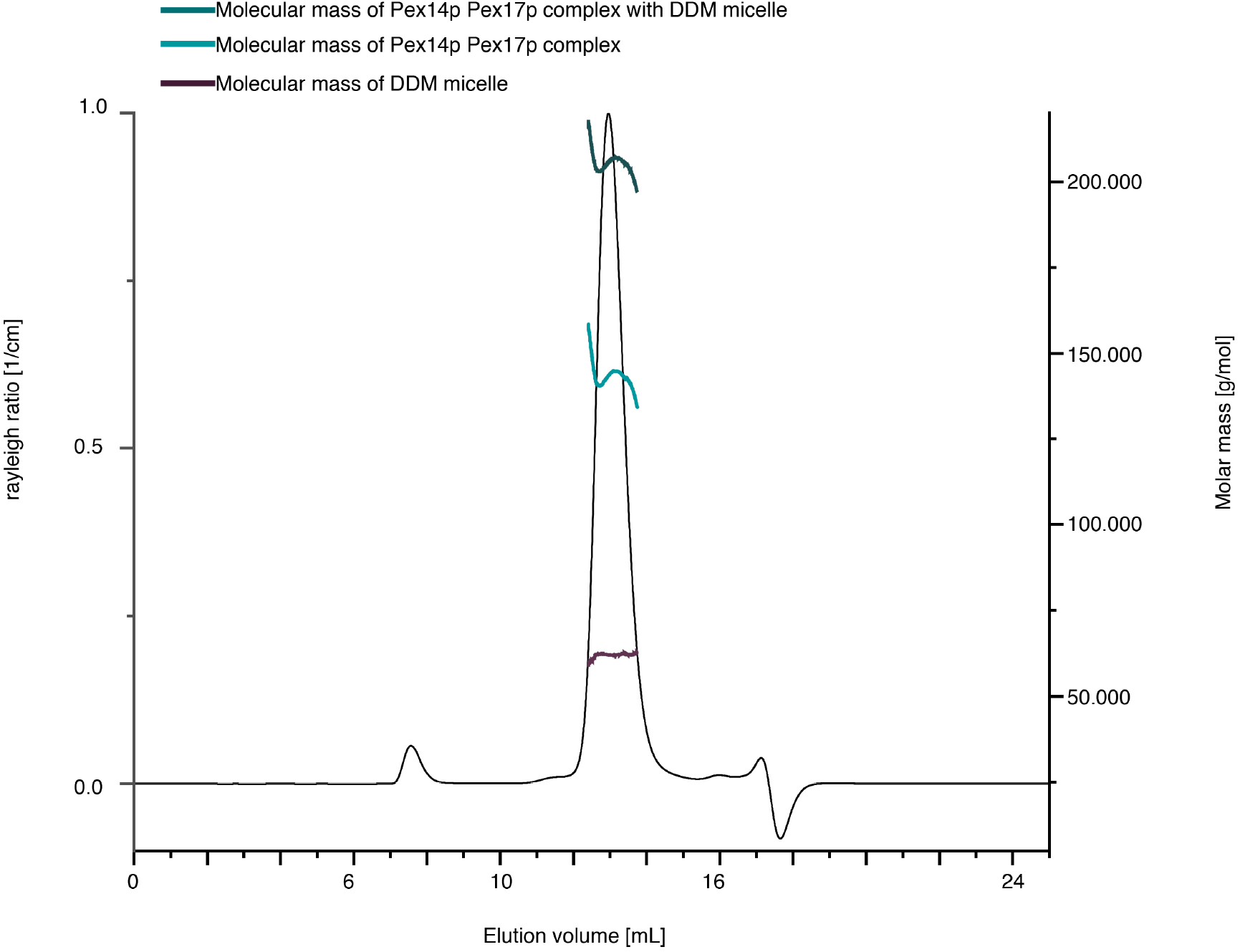
Chromatogram of light scattering Rayleigh ratio. The molar masses were calculated using the protein conjugate analysis. Superose 6 increase 10/300 column was pre equilibrated with buffer 50 mM Tris pH 8.0, 150 mM NaCl and 0.1% DDM. The protein molecular weight in the SEC peak ist ^~^200 kDA, and the associated detergent micelle of DDM is 62 kDA, respectively. This results in a molecular weight of 143 kDA for the Pex14p/Pex17p complex.

**Supplementary Figure 5.**
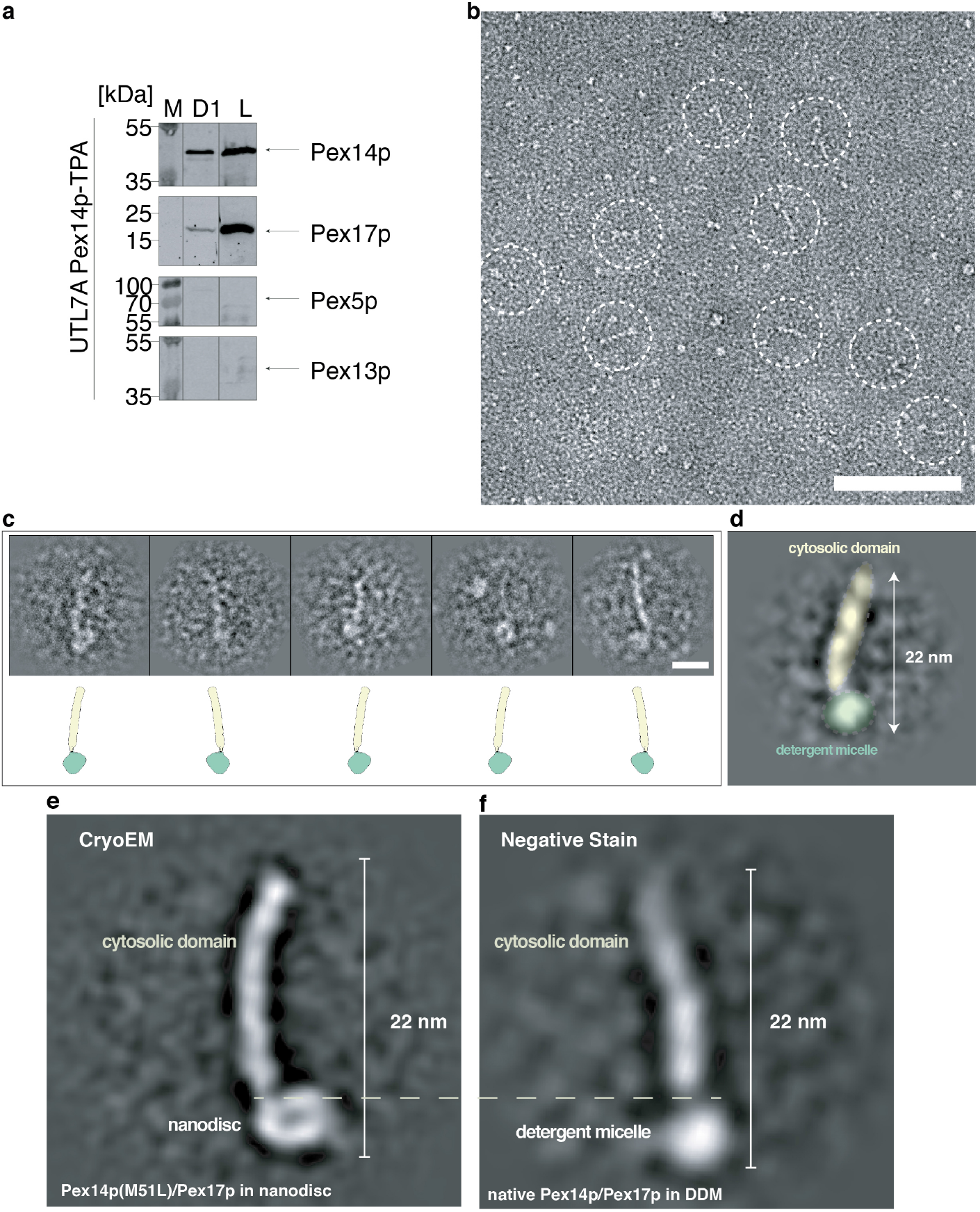
Negative stain EM analysis of the native Pex14p/Pex17p complex. **a)** Native complexes were isolated from Triton X-100 solubilized whole cell membranes from oleic acid induced cells expressing a Pex14p-TPA. Complexes were eluted from IgG-Sepharose by TEV protease cleavage. The obtained eluate was further analyzed by size-exclusion chromatography and subjected to immunoblot analysis with antibodies as indicated. The isolated native Pex14p/Pex17p complex in fraction D1 was subsequently analyzed with negative stain EM **b)** Representative negative stain micrograph of fraction D1. White circles indicate typical Pex14p/17p rod-like particles. **c)** Representative single particles. Scale bar 10 nm **d)** Representative reference-free class average of the native Pex14p/17p complex. Densities of the detergent micelle (green) and the cytosolic domains (yellow) are highlighted. **e-f)** Side-by-side comparison between reference-free class averages of the recombinantly expressed Pex14p(M51L)/His_6_-Pex17p in nanodisc (cryoEM) and the native Pex14p/Pex17p complex in DDM (negative stain).

**Supplementary Figure 6.**
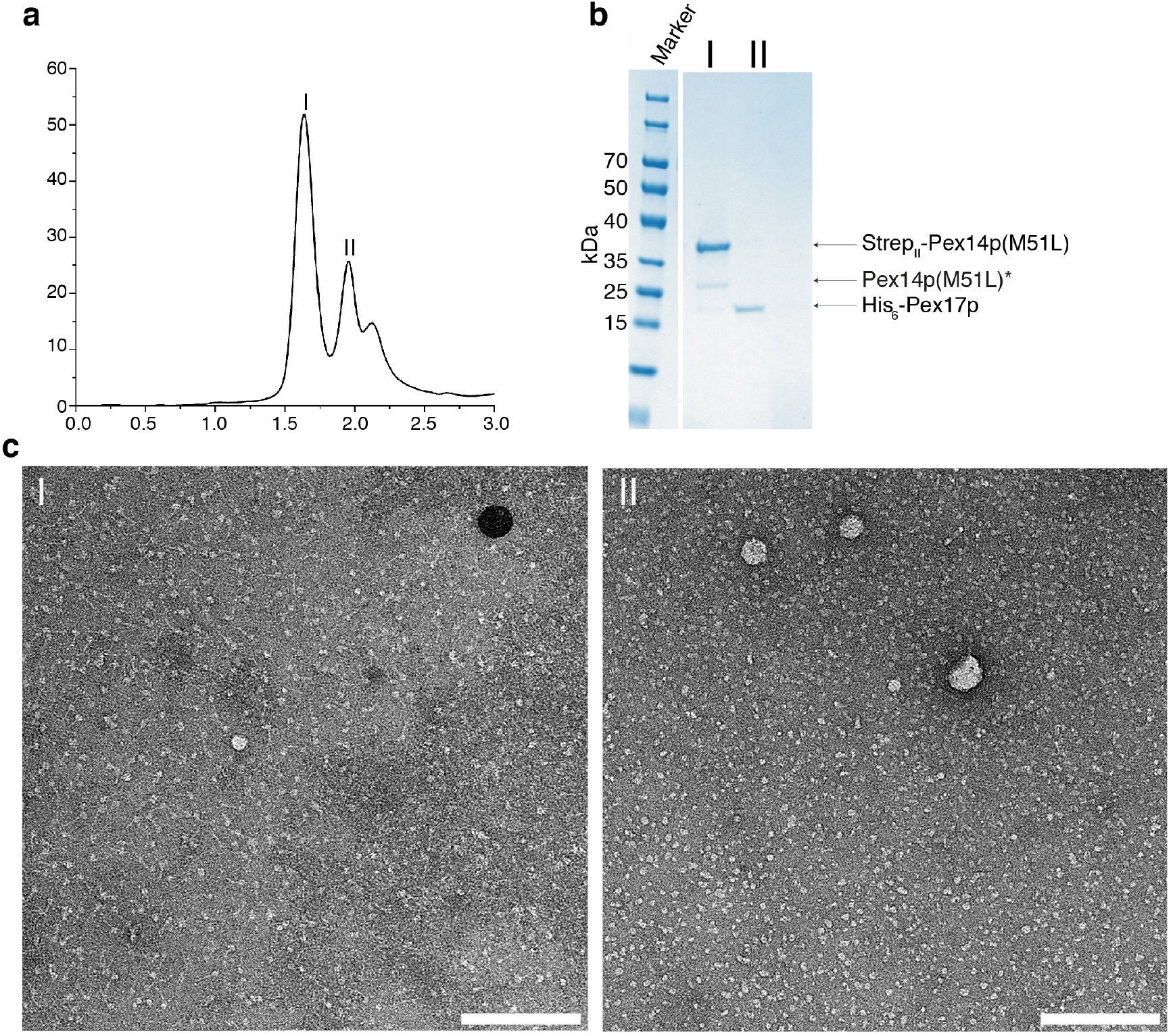
Pex14p/Pex17p dissociates after exchange of DDM with amphipols A8-35. **(a)** Size exclusion chromatography profile of Pex14pPex17p in amphipols. Peak I and II correspond to Pex14p and Pex17p, respectively. **(b)** SDS-PAGE of the peak fractions I and II after size exclusion chromatography. **(c)** Representative negative stain electron micrographs of the Pex14p (left panel) and Pex17p (right panel) fractions. Scale bar, 100 nm.

**Supplementary Figure 7.**
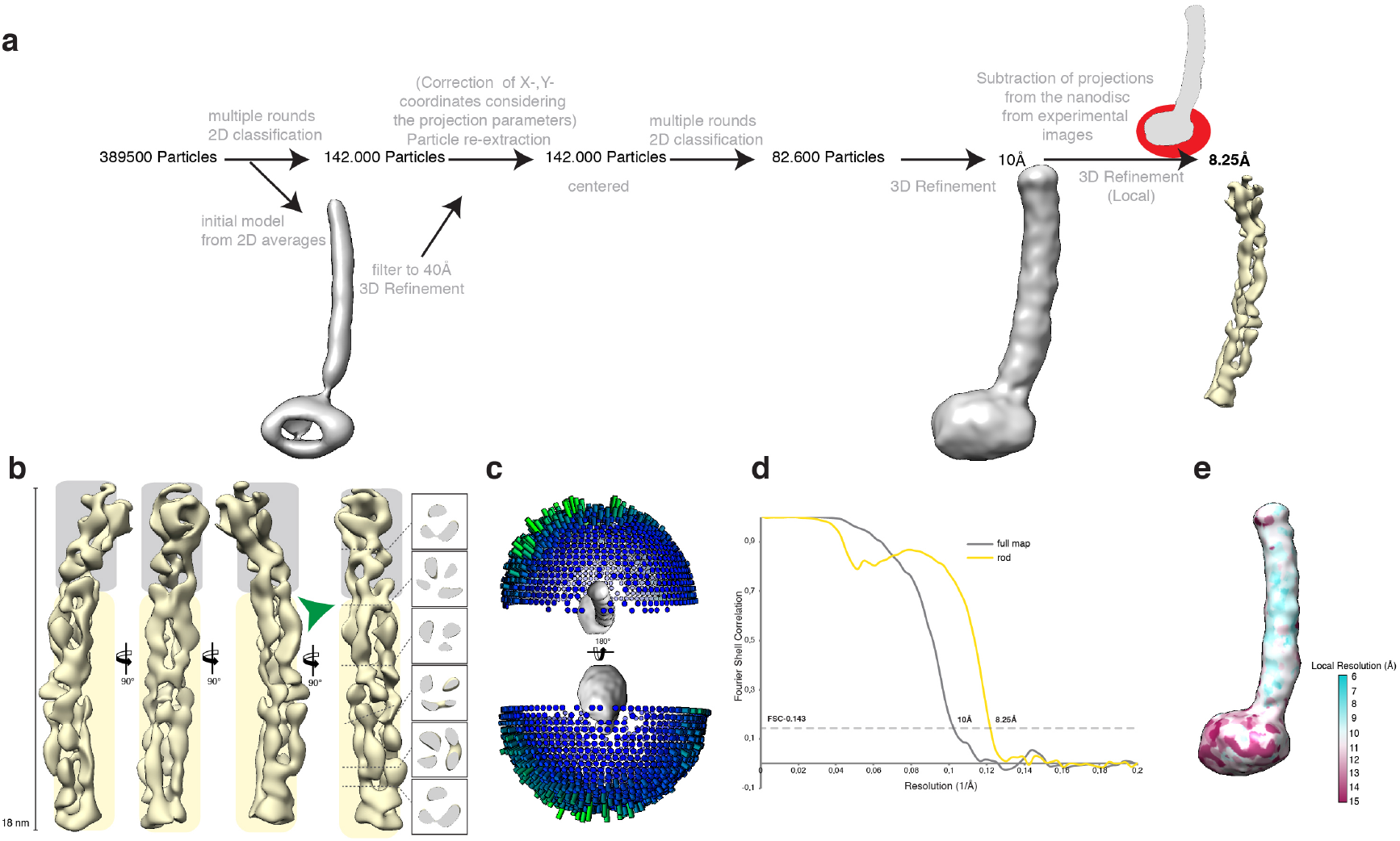
CryoEM analysis of Pex14p/Pex17p. **(a)** Single particle processing workflow for Pex14p/Pex17p structure determination. **(b)** Resulting map from focused 3D local refinement on the rod region after signal subtraction of the nanodisc. Insets show slices through the rod density at different heights. The two structural regions of the rod are highlighted in gray and yellow. The green arrowhead indicates the kinked connection between these two structural regions. **(c)** Orientation distribution of the particles used in the final refinement round prior signal subtraction. **(d)** Fourier Shell Correlation (FSC) curves between two independently refined half maps derived after standard refinement of the complete dataset and after signal subtraction of the nanodisc projections from the experimental images and subsequent local refinement (see, a). **(e)** Local resolution map of the Pex14p/Pex17p complex prior signal subtraction.

**Supplementary Figure 8.**
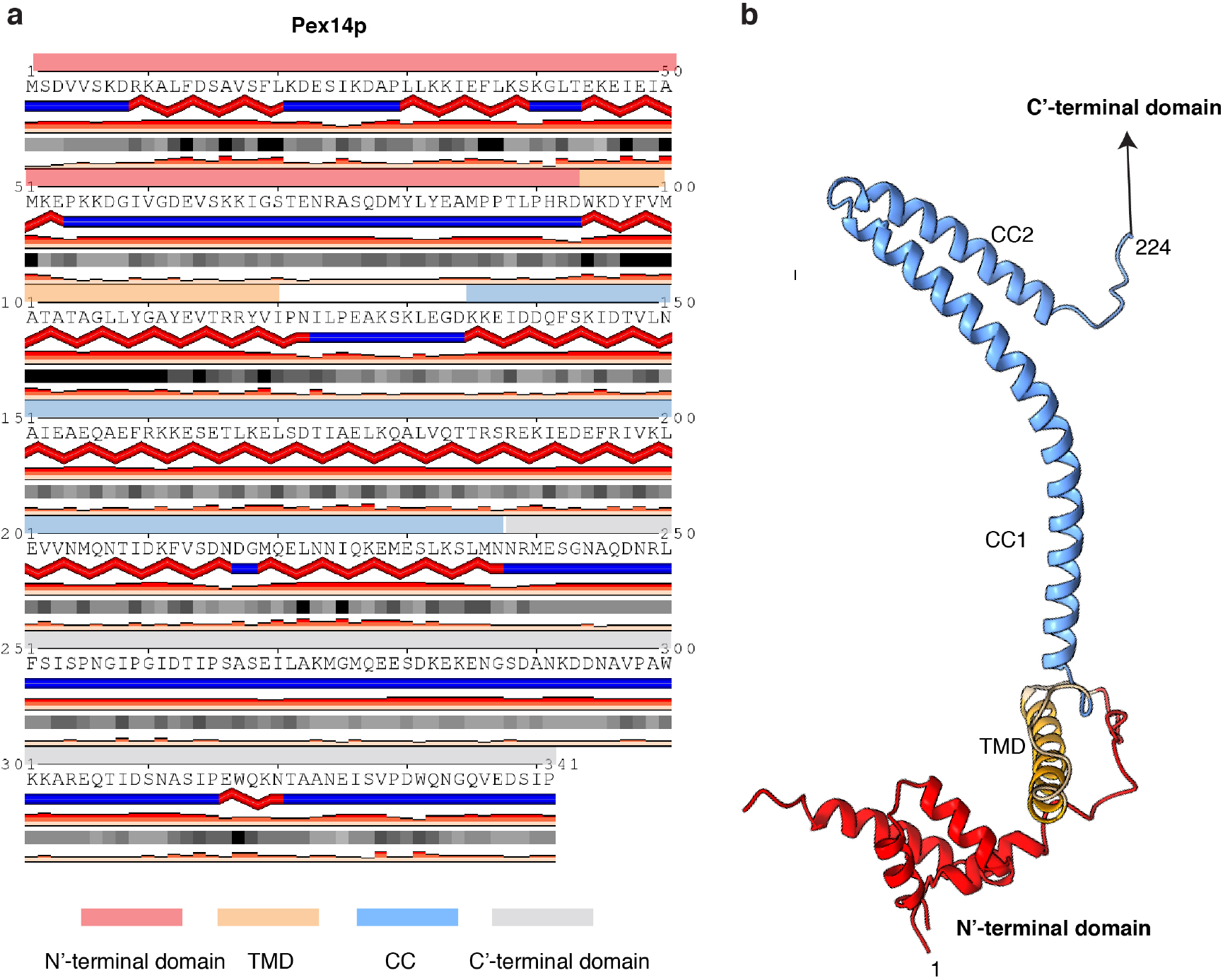
Structure prediction for the Pex14p subunit. **(a)** Sequence-based secondary structure prediction for Pex14p. Red bands represent α-helices. Blue-bands represent regions without predicted secondary structure. The subunit can be roughly divided into four domains (see also Figure 1a): N-terminal α-helical domain (highlighted in red), transmembrane domain (TMD; highlighted in brown), coiled-coil domain (highlighted in light blue) and the unstructured C-terminal domain (highlighted in gray) **(b)** Three-dimensional structure of Pex14p as predicted by RaptorX (*52*) (coloring similar to (a)).

**Supplementary Figure 9.**
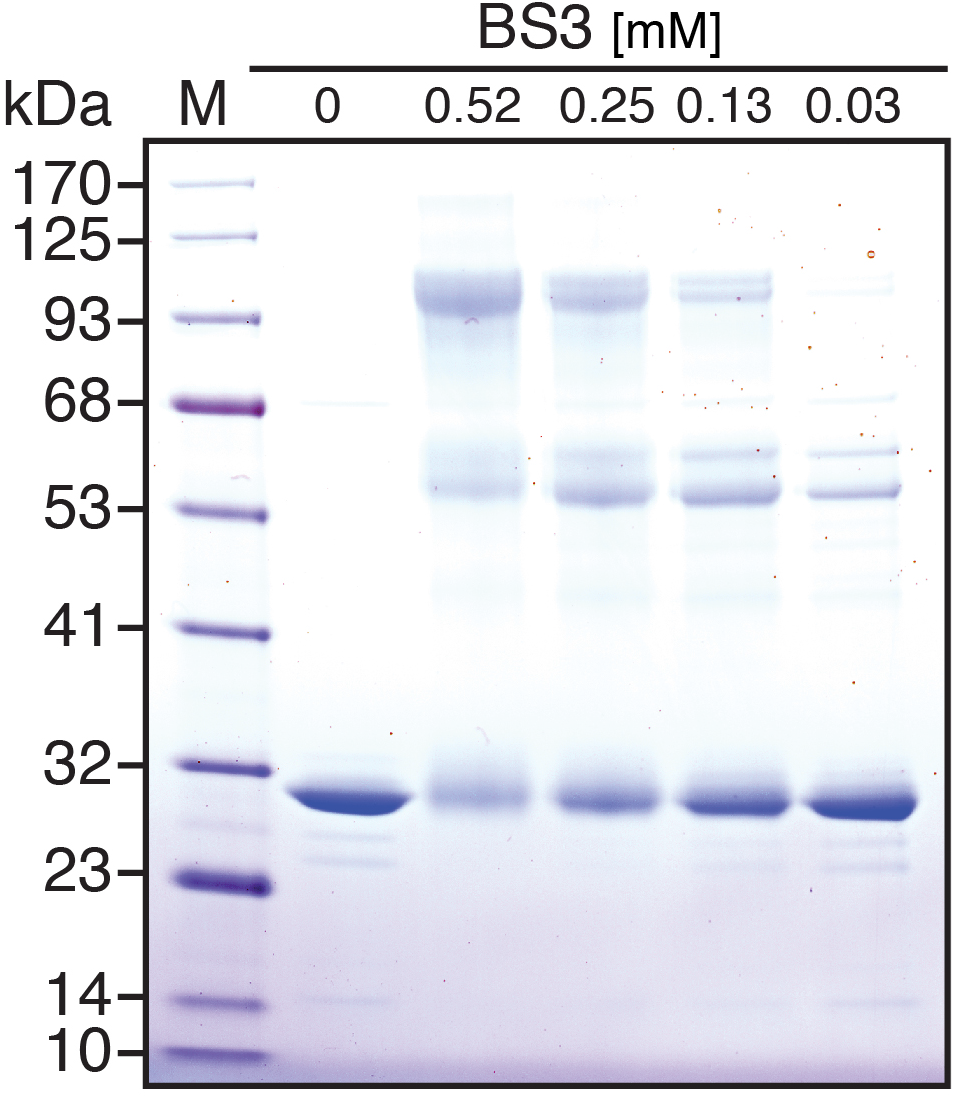
*In vitro* crosslinking leads to trimers of Pex14p_119-341_. Recombinant Pex14p_119-341_ was cross-linked using BS3 at increasing concentration as indicated and separated by SDS-PAGE followed by Coomassie staining.

**Supplementary Figure 10.**
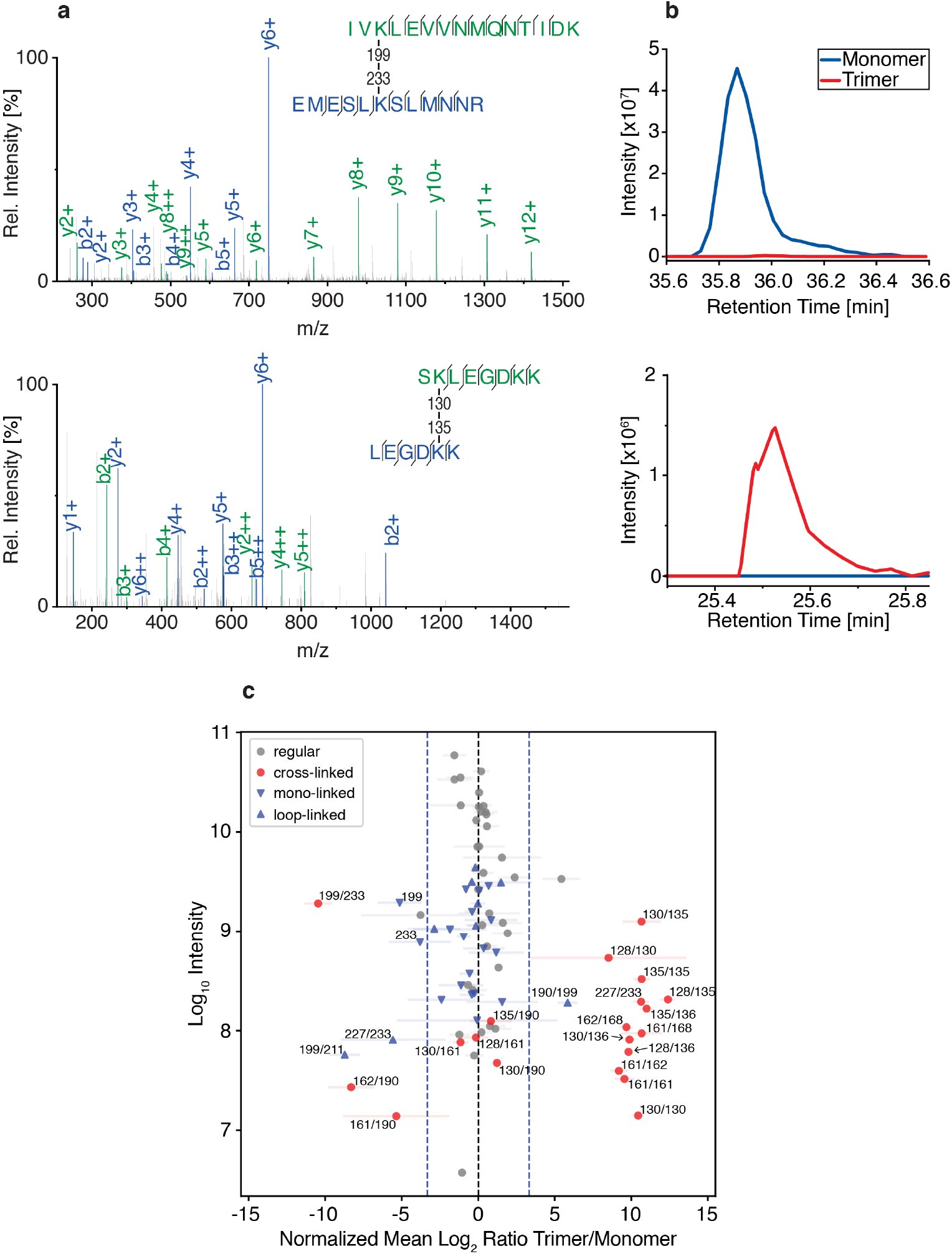
Intra- and intermolecular cross-links of Pex14p homooligomers. Following chemical crosslinking with BS3 and SDS-PAGE analysis, monomeric, dimeric and trimeric Pex14p_119-341_ forms (see Figure 4a) were subjected to in-gel digestion with trypsin and analyzed by LC-MS/MS. **(a)** MS/MS spectrum identifying a monomer-specific Pex14p intraprotein crosslinked peptide pair (top) and an oligomer-specific interprotein cross-linked peptide pair (bottom). Ions matched to masses of b- and y-ions (see inset) of the longer (green) and shorter (blue) peptides are marked. **(b)** Representative ion chromatograms showing the elution profiles of the corresponding monomer- (top) and trimer-specific (bottom) cross-linked peptide pairs. **(c)** Quantitative LC-MS analysis of cross-linked amino acid pairs (red dots), regular peptides (gray dots) and mono- (blue triangles down) or loop-linked amino acids (blue triangles up). Normalized log_2_ trimer-to-monomer intensity ratios were computed and plotted against their log_10_ intensities. Identified linkages are annotated with their sequence positions. Blue vertical lines indicate a 10-fold enrichment either in the trimer or monomer sample.

**Supplementary Figure 11.**
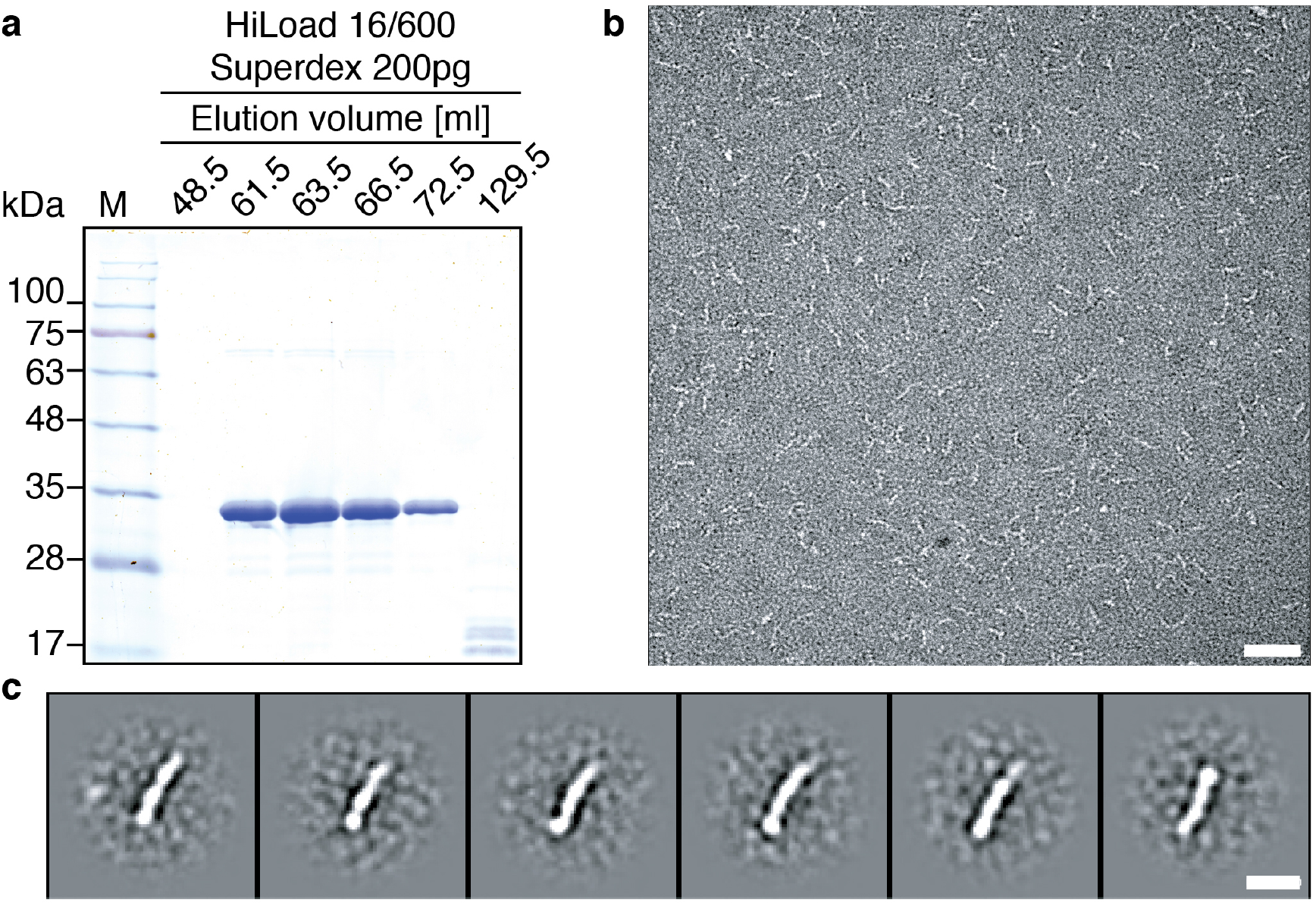
Negative stain EM analysis of the homotrimeric Pex14p_119-341_ variant. **a)** Coomassie stained SDS-Page of the Pex14p_119-341_ variant after purification using a HiLoad 16/600 Superdex 200. **b)** Representative EM micrograph with **c)** corresponding classes, scale bars 50 nm and 10 nm, respectively.

**Video 1. Flexibility of Pex14p/Pex17p in 2D.** The arrow indicates flexible densities below the nanodisc.

**Video 2. CryoEM map of the cytosolic rod.** The cryoEM map of the cytosolic domains obtained after signal subtraction of the nanodisc density from the experimental images and subsequent focused 3D local refinement is rotated in order to show the overall structure. α-helixes are fitted within rod-like densities. α-helices of the coiled-coil domain 1 of Pex14p are shown in blue (different shade of blue for each Pex14p subunit) and the helices of coiled-coil domain 1 and 2 of Pex17p are shown in yellow.

**Video 3. 3D segmentation of a Pex14p/Pex17p proteoliposome.**

